# An immunobiliary single-cell atlas resolves crosstalk of type 2 cDCs and γδT cells in cholangitis

**DOI:** 10.1101/2025.07.10.664083

**Authors:** Stefan Thomann, Helene Hemmer, Ankit Agrawal, Sukanya Basu, Judith Schaf, Sagar, Fabian Imdahl, Tanja Poth, Marcell Tóth, Christina E. Zielinski, Tobias Poch, Jenny Krause, Andreas Rosenwald, Katja Breitkopf-Heinlein, Nuh Rahbari, Dominic Grün

## Abstract

**Background and aims:** The immunobiliary niche serves as a reservoir of tissue-resident immune cells, yet the role of unconventional T cells during cholangitis remains poorly understood. Here, we connect cell state dynamics of type 2 conventional dendritic cells (cDC2) in cholangitis with site-specific *γδ*T17 responses in liver and draining lymph nodes (LN).

**Methods:** The 0.1% Diethoxycarbonyl-1,4-Dihydrocollidine (DDC) diet was used to generate an immunobiliary DC- and *γδ*T-enriched mouse liver and LN single-cell RNA-sequencing (scRNA-seq) atlas covering temporal disease dynamics, resolution and cDC2B depletion. cDC2 trajectories were inferred using VarID2 and *γδ*T cell transcription factor (TF) regulon activity was predicted by SCENIC. The human biliary niche was resolved by integrating human liver scRNA-seq data with available spatial transcriptomics data. Functional studies were conducted using Tcrd knockout, Il17a/f knockout and Tcrd reporter mice.

**Results:** A disease trajectory of Mgl2^+^ cDC2B was identified connecting DC maturation, homing and the expression of Il17-inducing genes. Disease progression was associated with numeric exhaustion of mature cDC2B and recruitment of DC precursors. *γδ*T cells were the main Il17 producers and were subtyped into Il17a^high^ *Scart1*^+^ V*γ*6^+^ and Il17^low^ *Scart2*^+^ V*γ*4^+^ populations exhibiting cDC2-directed communication and divergent TF regulon activity. Spatial proximity and conserved molecular interactions of cDC2 and *γδ*T cells were confirmed in human cholangitis. Il17-deficiency resulted in reduced liver fibrosis in mice, while cDC2B depletion attenuated *γδ*T17 cell states.

**Conclusions:** In cholangitis, a profibrogenic function of *γδ*T cells is contingent on the induction by peribiliary cDC2B, thereby highlighting relevant disease determinants within the immunobiliary and liver-draining LN niche.

**Impact and Implications:** The immunobiliary niche contains rare immune cells such as conventional dendritic cells and unconventional T cells, however, the function of these cell types in liver inflammation remains poorly understood. Mirroring human biliary diseases such as primary sclerosing cholangitis, we induced experimental cholangitis in a mouse model to generate a site-specific single-cell sequencing atlas resource resolving underexplored cell populations. Our data capture a profibrotic hepatic disease state trajectory of Mgl2^+^ cDC2B inducing a *γδ*T cell-specific Il17-response, which is attenuated upon cDC2B depletion and in DC precursors, which are characterized by reduced genomic accessibility of *γδ*T cell-interacting genes.

These results highlight the importance of portal niche residing underexplored immune cell populations and the necessity to further resolve immunobiliary niche responses and crosstalk in inflammatory settings such as in cholangitis.

**Graphical Abstract:** 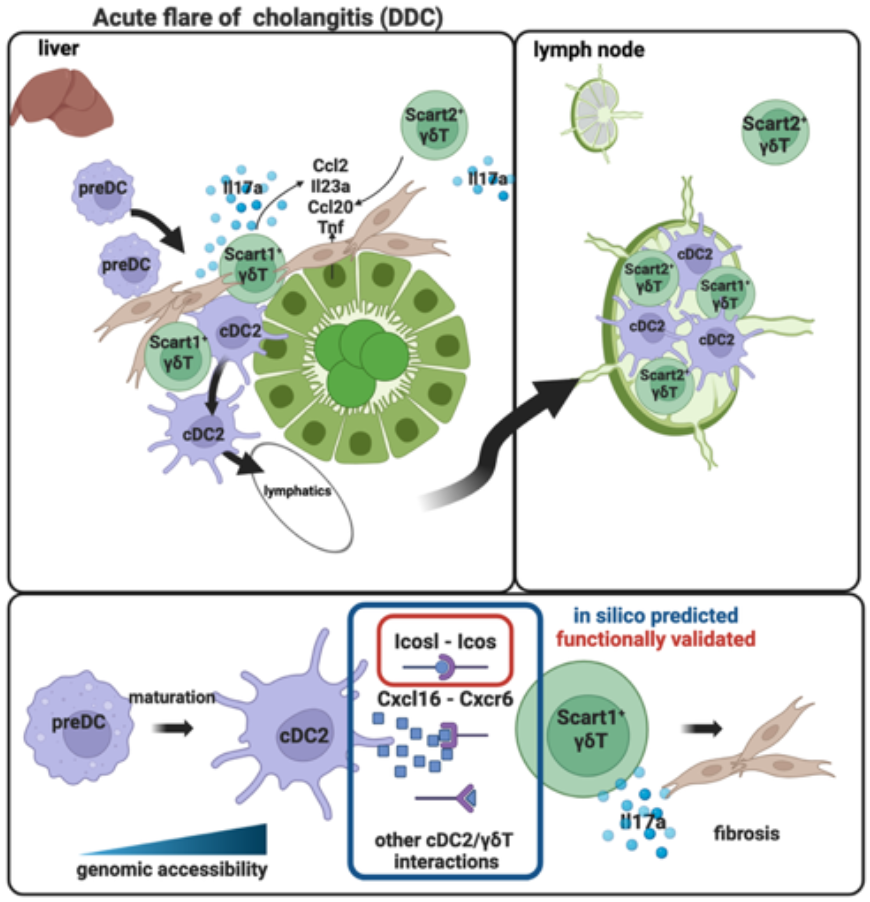

## Introduction

Primary sclerosing cholangitis (PSC) is a multifactorial autoimmune-related disease with rising occurrence in the US and EU [1, 2], and the disease course ultimately requires liver transplantation. While there are different subtypes of PSC [3], and PSC is frequently linked to other chronic inflammatory diseases, the common histological features include periductal fibrosis, proliferation of reactive bile ducts known as ductular reaction (DR), bile duct stenosis, dilatation and rarefication leading to biliary cirrhosis and dysfunctional myeloid responses [4, 5]. Clinical presentation of PSC and its chronic disease activity may include intermittent episodes of acute cholangitis [6] and involve dysfunctional cDC2 and T cell responses contributing to disease severity [7, 8], yet the role of unconventional T cells (UTC) and their potential crosstalk with cDC2 remains to be elucidated.

In the hepatic cDC2 niche, continuous reconstitution by circulating cells and the DC life-cycle have been characterized in steady-state and disease [9, 10], and follow the steps of liver sinusoid attachment, transmigration to the space of Dissé and migration towards the portal field stroma [11]. The portal microenvironment may furthermore instruct DCs towards a more anti-inflammatory phenotype [12], however, a significant fraction of the liver cDC2 pool has been characterized as a proinflammatory cDC2B phenotype [13] and the behavior of recently recruited DCs versus niche-adapted cells remains challenging to resolve in inflammation and resolution.

In this study, we explore the tissue niche, differentiation dynamics, and functional roles of cDC2 during cholangitis in the 0.1% 3,5-Diethoxycarbonyl-1,4-Dihydrocollidine Diet (DDC)-diet mouse model [14]. By administering DDC for durations of 3 – 25 days and following inflammatory resolution for 7 days, we established a high-resolution temporal and multimodal single-cell RNA-sequencing (scRNA-seq) atlas covering the cellular dynamics that shape the immunobiliary niche and the liver draining lymph nodes (LN). Combined with spatial analysis, this resource provides novel insights into differentiation dynamics of cDC2 and their heterocellular interactions and complements available spatiotemporal DDC/PSC data of more abundant cell types [5, 15]. Our data reveal crosstalk of cDC2 with *γδ*T cells governing the inflammatory response, which we confirm in other acute models of liver injury and functionally validate *in vivo* with the help of diphtheria toxin receptor (DTR)-based depletion of cDC2B using the CD301b-DTR (Mgl2-DTR) model [16]. Together, our work highlights new site-specific and - independent avenues of cDC2B – *γδ*T cell interactions in cholangitis.

## Materials and Methods

### Mouse Models

All animal experiments were authorized by the German Regional Council of Baden-Württemberg and Bavaria (Regierungspräsidium Freiburg and Regierung Unterfranken, reference numbers G20/164, 2-1554). Mice were bred in a 12-hour light/dark cycle with free access to water and food and kept under defined pathogen free conditions. C57BL/6J were bought from Janvier (Saint-Berthevin Cedex, France). Il17a/f-KO [17] and Tcrdtm1 mice [18] were obtained from Wolfgang Kastenmüller. Tcrd-GDL mice [19] were obtained from Martin Väth. Mgl2-DTR [16] were obtained from Kristin Hogquist with permission from Akiko Iwasaki. Chow supplemented with 0.1% 3,5-Diethoxycarbonyl-1,4-Dihydrocollidine (DDC) or chow deficient in choline was purchased at ssniff (Soest, Germany) [14]. For CDE diet experiments, mice received a chow deficient in choline and 0.05% ethionine-supplemented drinking water.

For scRNA-seq experiments male mice were used. For FACS analysis male mice were used with the exception of the cDC1/2 and cDC2A/B quantifications, where female mice were used.

### Isolation of cells, FACS and GEM generation

Primary liver cells were isolated from 6 – 12-week-old C57BL/6J mice as previously described [20]. Prior to FACS sorting, cells of interest were magnetically enriched using magnetic separator and antibody-or Streptavidin-conjugated microbeads (Miltenyi Biotec GmbH, Bergisch Gladbach, Germany). Isolates of hepatic stellate cells (HSCs) were obtained as described elsewhere [21]. Mouse hepatoduodenal and celiac lymph nodes [22] were excised, and digested in 2mg/ml Collagenase D for 30 min at 37° C. Live/dead exclusion in FACS was performed using Zombie-NIR fixable viability dye (Biolegend, San Diego, USA) in combination with antibodies (supplemental methods table) to sort the cell populations at equal ratios using a FACS Aria III cell sorter. Information regarding the composition of individual cell populations in each dataset can be found in the supplemental information. GEM generation and library preparation was performed according to the manufacturer’s protocol (10x Genomics, Pleasanton, USA). Quality control of libraries was performed using Agilent Bioanalyser High Sensitivity DNA Chips. Single and dual index libraries were sequenced on a NovaSeq S1 or NextSeq 2000 with a read-depth of 40 000 reads/cell. FASTQ files were mapped to a reference using kallisto [23] and the GRCm38.99 cDNA reference (DDC atlas data). CellRanger was used for mapping of transcriptome, antibody-derived tags and human data. Combined scRNA- and scATAC-seq (10x Multiome) was performed as described in the demonstrated protocol for nuclei isolation for single cell Multiome ATAC + Gene Expression Sequencing (CG000365.Rev C). Digitonin treatment was performed for 210 s, nuclei suspensions washed, quantified and loaded at a concentration between 3000 – 8000 nuclei/µl to target a recovery of 10,000 nuclei. Before proceeding to GEM generation, a transposition reaction was performed on the isolated nuclei, where transposase enzymes fragmented accessible chromatin regions and inserted sequencing adapters. Nuclei were then loaded into the 10x Genomics Chromium Controller on a Chip J, where they were partitioned into Gel Bead-In Emulsions (GEMs), enabling the simultaneous capture of gene expression (GEX) and chromatin accessibility data from the same nuclei. Within the GEMs, reverse transcription converted mRNA into barcoded cDNA, unique to each nucleus. Following GEM breaking, the cDNA was recovered and amplified, alongside the tagmented DNA fragments from the ATAC reaction. Both the GEX and ATAC libraries were then purified, amplified, and quantified separately. Finally, these libraries were sequenced on separate flow cells, as the GEX and ATAC libraries require different sequencing parameters. Transposition and Chip loading for Multiome experiments were conducted at the Single Cell Center of the University of Würzburg.

### Bioinformatic analysis of scRNA-seq data

For the initial temporal DDC dataset (Figure 1), a threshold of 1500 transcripts for mouse and 1000 transcripts for human datasets was used. VarID2 [24] was run with the following parameters: mintotal = 1500, minexpr = 5, minnumber = 5, ccor =0.4. FGenes was used to remove genes without gene symbol, mitochondrial and ribosomal genes. The pruneKnn function was run with the parameters: large=TRUE, regNB=TRUE, knn=25 and Leiden clustering performed.

**Figure 1.**
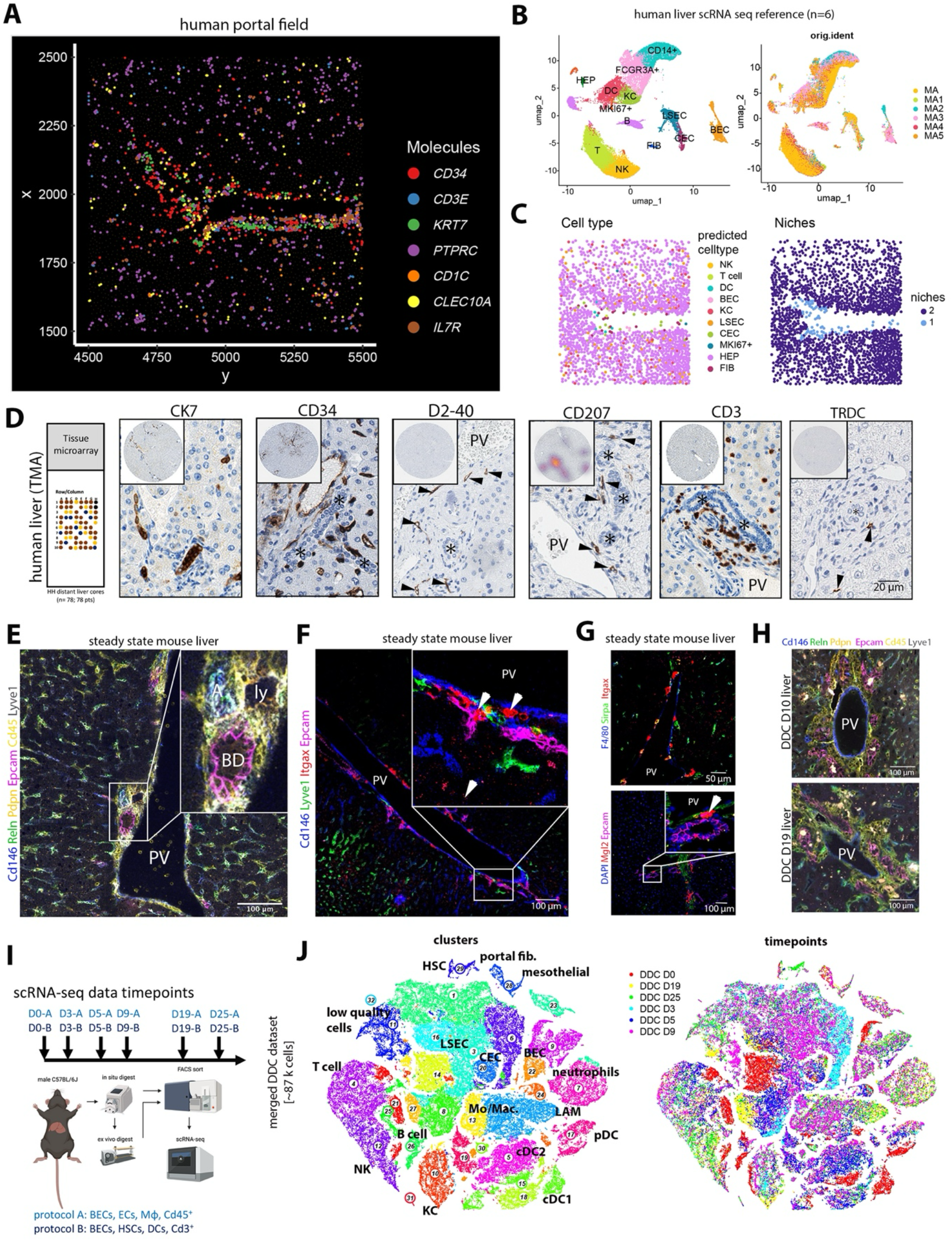
Immunobiliary microenvironment in human and mouse. **(A)** Spatial transcriptomics data subset of a human portal field, displaying expression of 7 RNA molecules of interest in the immunobiliary niche. **(B)** UMAP representation of a human control liver scRNA-seq reference (32,901 cells from 6 patients) highlighting major cell types of the portal niche (left) and individual samples (right). **(C)** Spatial map highlighting cell type composition predicted by label transfer (left) and inferred niche domains (right). **(D)** Immunohistochemical stainings of a human TMA containing pseudonormal liver tissues including CK7^+^, CD34^+^, PDPN^+^ (clone D2-40), CD207^+^, CD3^+^, TRDC^+^ cells (arrowheads). Asterisks demarcate bile ducts adjacent to portal veins (PV). Small magnification displays TMA core, and large magnification portal region containing cell types of interest. Density map encodes local cell concentration and indicates local enrichment of CD207^+^ cells within portal fields. **(E)** IF of major cell type markers of immunobiliary niche on steady state mouse liver tissue. High magnification displays the presence of Cd45^+^ immune cells close to PV, hepatic artery (A), bile duct (BD) and lymphatic vessels (ly) (n=3 mice). **(F)** IF of steady state mouse liver portal fields visualizes close spatial proximity of Epcam^+^ BECs with Itgax^+^ cells (n= 3 mice). **(G)** IF showing Itgax^+^ Sirpa^+^ cells (top) and close proximity of Mgl2^+^ cDC2B and Epcam^+^ BECs (bottom) in steady state liver tissue (n= 3 mice). **(H)** IF of major cell type markers on DDC D10 and D19 mouse liver tissue visualizes expansion of the immunobiliary niche. (n=4 mice/group). **(I)** Scheme of DDC scRNA-seq disease atlas containing a control (D0) and 5 disease timepoints (3 – 25 days). Two isolation protocols with different enrichment of cell types were used per timepoint. **(J)** tSNE representation of the complete DDC atlas data (∼87k cells) highlighting cell type annotation (left) and individual timepoints (right).

For the analysis of Xenium data, we used publicly available liver control data (see methods table). Xenium data were processed within Seurat and principal components, UMAP, community detection and clustering information obtained. For the inference of human suspension-based liver scRNA-seq data, we used robust cell type decomposition (RCTD) as part of the spacexr package [25]. For the identification of common cellular neighborhoods, we used Seurat’s “BuildNicheAssay”, which identifies similarities based on k-means clustering of neighborhood data (neighbors.k = 15). Transition probabilities within cDC2 subsets were calculated using the plotTrProbs function of VarID2 [24]. Pseudotemporal ordering of self-organizing maps (SOMs) was performed using FateID [26]. Signac [27] was used for the analysis of merged D0, DDC D5 Multiome data. RNA and ATAC data were tested for quality metrics and clustered based on information within each modality. Predicted gene activities levels were compared to RNA expression within transcriptome-based dimensionality reduced space.

Seurat’s ability was used to integrate multimodal data including antibody-derived tag (ADT) data [28]. ADT data was transformed using centered log ratio transformation (CLR) and ADT data displayed within the gene expression space. SCENIC [29] was used to infer transcription factor regulon activity. Regulon activity per cell type was visualized in a heatmap and regulon specificity scores (rss) calculated using the inbuild calcRSS function. Pathway enrichment analysis was performed using ReactomePA [30]. cDC2A/B gene signatures were derived by curation from [13]. Residency and circulatory gene signatures [31, 32] were derived from [33]. V*γ*4 and V*γ*6 gene signatures were calculated by including the top 50 differentially expressed genes within a V*γ*4 cluster (1) and V*γ*6 cluster (9) of a TCR repertoire information containing reference [33]. For Niche Covariation (NiCo) analysis [34] of the human liver Xenium dataset, we utilized NiCo’s annotation module with spatial guide clustering resolution 0.5.

More detailed information about the materials and methods are stated in the supplemental information.

## Results

### Cell type composition of the immunobiliary niche in human and mouse

To determine spatial immune cell type localization in human steady state liver, we first explored publicly available spatial transcriptomics (ST) data (Methods) and found enrichment of *CD1C* and *CD3E* transcripts near large vessels and biliary tracts (Fig. 1A). For cell type annotation of this high-resolution ST reference, we generated a human pseudonormal liver scRNA-seq reference dataset of the immunobiliary niche comprising ∼33k cells from 6 patients, by FACS-sorting biliary epithelial cells (BECs) and immune cell populations (Fig. 1B, S1A-C). We performed spatial cell type annotation by running RCTD [25], which defined DCs, continuous endothelial cells (CECs), BECs and T cells to be part of the immunobiliary niche (Fig. 1C). Topological niches were predicted by Seurat’s *BuildNicheAssay* and confirmed the existence of a portal immunobiliary (“1”) and lobular niche (“2”; Fig. 1C), providing evidence for the immunobiliary niche as a T cell- and DC-containing microenvironment. These findings were confirmed using pseudonormal human liver tissues derived from a characterized tissue microarray (TMA) [35] by immunohistochemical detection of CK7^+^ BECs, CD34^+^ CECs, PDPN^+^ lymphatics (clone D2-40), CD207^+^ cDC2, CD3^+^ T cells and TRDC^+^ cells (Fig. 1D). The immunobiliary niche was conserved in mouse, where multicolor immunofluorescence (IF) revealed co-localization of Epcam^+^ BECs with Cd146^+^ CECs and Cd45^+^ immune cells (Fig. 1E, S2A). Itgax^+^ cells were detected in the biliary niche of steady state mice and co-expressed CD172a (encoded by *Sirpa*) and peribiliary CD301b expression (encoded by *Mgl2*) was equally detected, likely representing cDC2 (Fig. 1F, G).

To locally perturb the immunobiliary niche, we made use of a xenobiotic diet, that is known to induce cholangitis and ductular reaction through supplementation of DDC (Fig. 1H) [14]. qPCR detected DDC-induced upregulation of biliary and myeloid marker genes suggested expansion of these populations (Fig. S2B). Physical interaction of CD45^+^ immune cells with Epcam^+^ bile ducts was validated by IF image co-localization, indicating a dynamic immunobiliary crosstalk upon ductular reaction (Fig. S2C-E). To unbiasedly investigate dynamics of gene expression across cell types residing in the immunobiliary niche, we performed scRNA-seq on liver tissue isolated at five different timepoints of DDC diet (day (D) 3, D5, D9, D19, and D25) and from control liver (D0) (Fig. 1I). Based on two independent isolation protocols (see methods), we were able to capture abundant cells of the liver immunobiliary niche, as well as rare liver immune cells such as cDCs (Fig. 1J, S2F-H, Suppl. Table 1).

### DDC diet induces small duct inflammatory disease

To dissect the BEC subset of our scRNA-seq atlas, we performed a separate clustering analysis of the two main cholangiocyte populations (clusters 9,22; 5,137 cells, Fig. 1J) resulting in 16 more granular clusters (Fig. 2A, S3A). Our data recapitulates two major subsets: *Dmbt1*^+^ *Muc4*^+^ *Sox17*^+^ BEC (cluster 2,6,11) and *Sox9*^+^ BEC (remaining clusters). Among *Sox9*^+^ BEC, clusters 4 and 8 showed an increased expression of chemotaxis- and adhesion-regulating genes (*Ccl2, Ccl20, Vcam1, Mif, Csf1, Tnf, Il23a*) (Fig. 2B, C), which are known cDC2 and T cell regulators. In particular, Il23 is key a cytokine for the induction of an Il17-response which is a known driver of liver inflammation and fibrosis [36]. Increased transcript abundance of *Tnf* and *Il23a* was confirmed within DDC tissue at D10 and D19 using qPCR (Fig. S3B).

**Figure 2.**
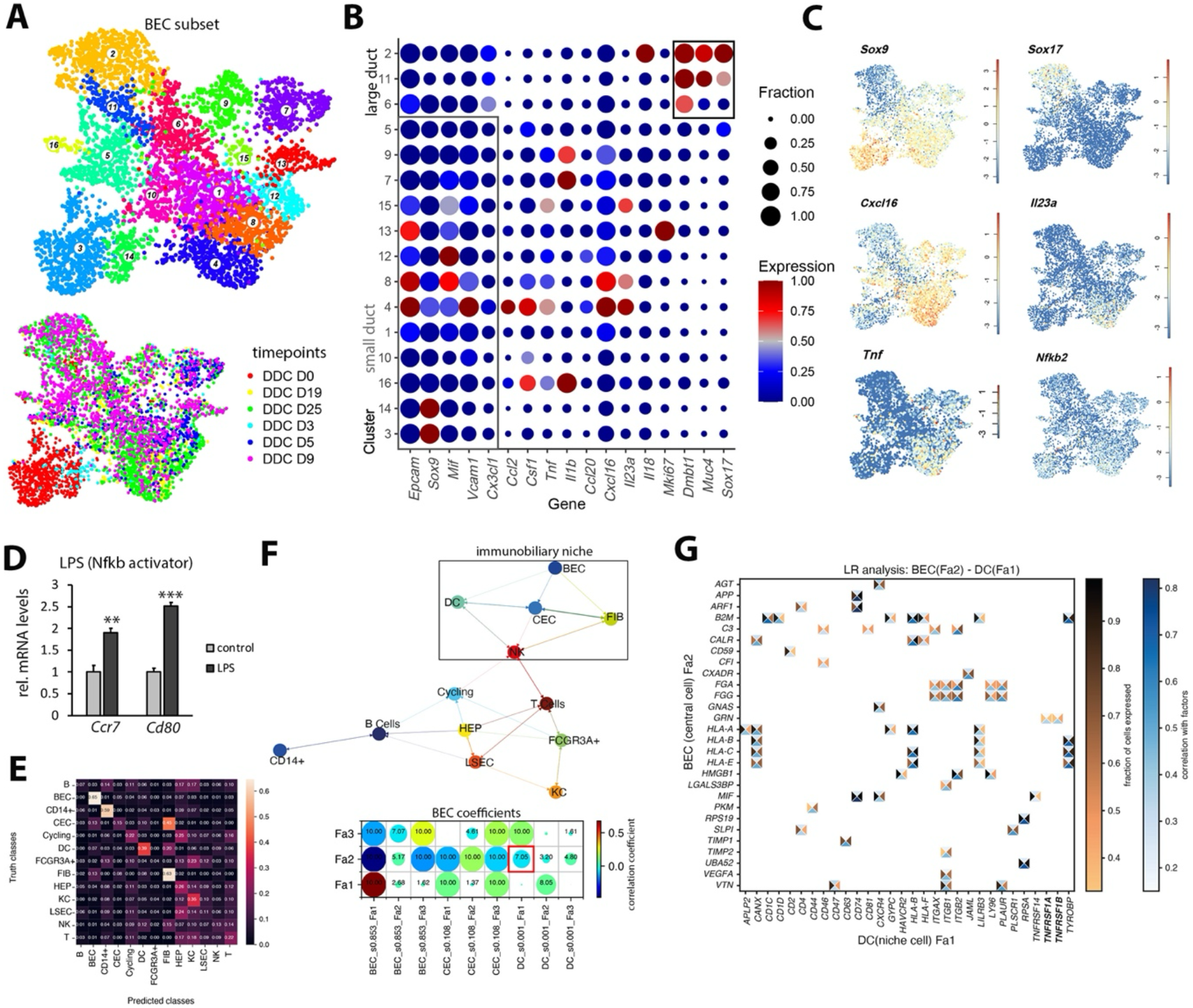
DDC induces ductular reaction and small duct inflammatory disease. **(A)** UMAP representation highlighting BEC sub-types (top) and individual timepoints (bottom). **(B)** Dotplot displaying BEC cluster-specific gene expression. Z score of the mean expression is color coded and fraction of cells expressing the gene is encoded by dot size. **(C)** UMAP of log-normalized expression of BEC and inflammatory markers. **(D)** Barplot displaying relative tissue mRNA levels of *Ccr7* and *Cd80* in cultivated liver DCs for untreated control and after LPS treatment. SD, standard deviation across technical replicates. Two-sided t-test (two independent experiments). Data of one experiment is shown. **p < 0.01, ***p < 0.001. **(E)** Confusion matrix displaying NiCo-based central cell identity prediction based on surrounding niche cell type frequencies within a human portal field (same as Fig. 1A). **(F)** Top: Spatial cell type interaction map derived by NiCo. Normalized logistic regression coefficient cutoff c = 0.07. Bottom: Regression coefficients between latent BEC factors (y-axis) and colocalized niche cell types (x-axis). Circle size encodes p-value (-log_10_ scale), circle color encodes ridge regression coefficients. **(G)** Ligand-receptor pairs correlated with covarying BEC Fa2 and DC Fa1 (cc, central cell; nc, niche cell). The rectangle’s north and south faces represent ligand and receptor correlation to the factors, while west and east faces represent the proportion of cells expressing the respective ligands or receptors. Gene correlation threshold 0.15 and gene expression threshold of 0.33 was applied. Ligands, y-axis; receptors, x-axis.

To assess the general nature of this pro-inflammatory early liver damage response, we revisited published data of a bile duct ligation (BDL) model and a chronic model of repetitive CCl4-induced liver damage [37]. Acute biliary injury (BDL) showed highly similar gene expression patterns within *Sox9*^+^ BECs, while chronic CCl4-induced damage did not lead to upregulation of *Tnf, Il23a, Cxcl16* and *Nfkb2* expression (Fig. S3C,D). Intersecting upregulated genes within our *Sox9*^+^ BEC DDC data (clusters 4,8 vs. 3,14) and *Sox9*^+^ BEC BDL data (clusters 3,4,7 vs. 1,2,9) revealed a common set of 128 genes (Fig. S3E) which were enriched for gene ontology annotations related to cell communication and inflammatory response (Fig. S3F). These findings highlight the small duct BEC as important cDC2/T cell niche regulators in acute liver damage, independent of the noxious background (DDC/BDL), where Tnf, as a Nfkb pathway inducer, may trigger a proinflammatory change within the cDC containing immunobiliary niche. To functionally test Nfkb-driven alterations in biliary niche-residing cDCs triggered by BEC-secreted Tnf, we treated cDCs isolated from control livers with a strong Nfkb pathway inducer (LPS) *in vitro* [38] and observed induction of the maturation markers *Ccr7* and *Cd80* in cDCs, indicating increased maturation dynamics (Fig. 2D).

To test the existence of an analogous Tnf-inducible Nfkb pathway activation node in the human liver immunobiliary niche within DCs, we made use of our human ST liver dataset and Niche Covariation (NiCo) analysis within a representative portal field data subset [34]. Cell type interaction prediction by NiCo recapitulated the immunobiliary niche and indicated a cellular localization of DCs within a defined cell type neighbourhood hosting BECs and CECs (Fig. 2E,F and S3G,H). Since in steady state we expected an inactivity of this communication node, we screened for covariation of BEC and DC gene programs represented by latent factors (Fa), and detected a negative covariation of BEC Fa2 and DC Fa1 (Fig. 2F,G). Correlating ligand-receptor pairs contained an interaction of BEC-derived *GRN* with DC-derived *TNFRSF1A/B*, an interaction mediating either pro-or anti-inflammatory functions dependent on the proteolytic cleavage of the gene product [39].

In conclusion, these results highlight small duct BEC-derived conserved and model-independent inflammatory gene expression patterns that may modify the surrounding myeloid niche.

### Disease stage-specific cell state transition in cDC2 reflects immature niche restoration

Based on their spatial proximity to the BEC compartment and their capacity to activate lymphocytes, we explored the temporal disease dynamics of DC in our DDC disease atlas. While DC subtype frequencies were approximately equally distributed in steady state, a frequency shift towards cDC2 predominance was observed starting from D3 (Fig. 3A). To confirm this shift by FACS analysis, we made use of a previously published gating strategy to discern cDC1 and cDC2 [40] (Fig. S4A). Increased cDC2 proportions at D5/6 and D10 after DDC diet were observed extending previous observations on frequency changes within the cDC1/2 compartment in short-term DDC-fed mice [7] (Fig. 3B). Cell state dynamics of cDC1/2 were comparably affected in a model of acute hepatotoxic injury and ductular reaction (CDE diet; Fig. S4B).

**Figure 3.**
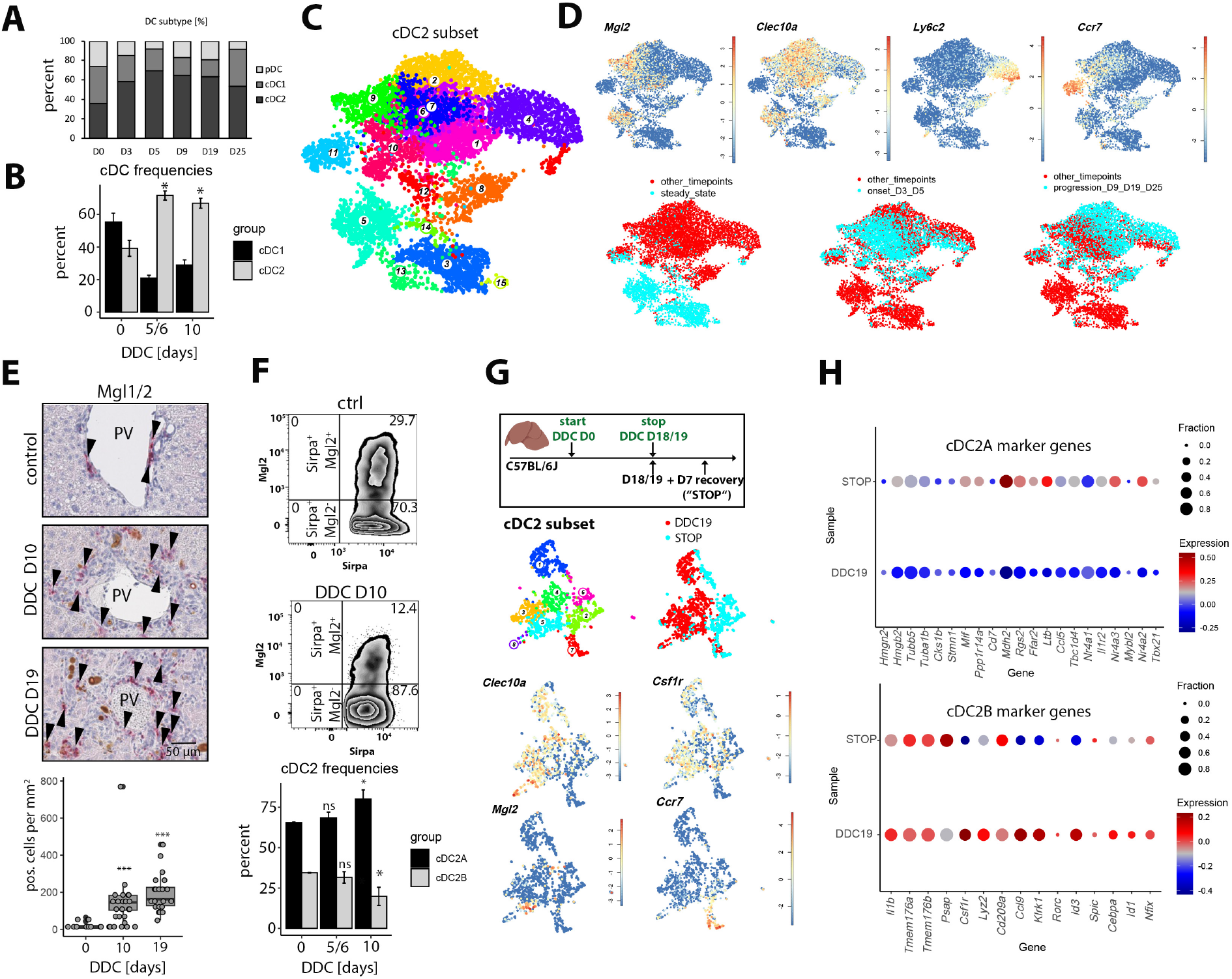
Disease stage-specific cell state transition in cDC2 reflects immature niche restoration. **(A)** Barplot displaying DC-subtype frequencies per timepoint in the DDC atlas data. **(B)** Barplot displaying cDC1/cDC2 FACS-based quantification at D0, DDC D5/6 and D10. One-sided Wilcoxon test (n=3-4 mice per group, two independent experiments) *p < 0.05. **(C)** UMAP representation of cDC2 subset (Cl.5,15 from Fig. 1J). **(D)** UMAP representation highlighting log-normalized expression of cDC2B marker genes (top) and timepoints of different disease stages (bottom). **(E)** Top: Immunohistochemical staining of Mgl1/2 in control and DDC D10/D19 liver tissues. Arrowheads highlight Mgl1/2^+^ cells. Bottom: Quantification of Mgl1/2^+^ cells across conditions. Every dot represents a tissue section that has been quantified. Wilcoxon rank sum test (n=3-4 mice/group), ***p ≤ 0.001. **(F)** FACS plots displaying hepatic LC^−^ Cd64^−^ MHC^+^ Itgax^+^ Sirpa^+^ cells (top) and quantification of relative fraction of Mgl2^+^ cDC2B (bottom). One-sided Wilcoxon test (two independent experiments, n=3-4 mice/group), *p ≤ 0.05. **(G)** Top: experimental design comprising a DDC D18/19 and a recovery (DDC18/19 + D7 recovery; “STOP”) timepoint. Bottom: UMAP representation of cDC2 subset, samples and log-normalized expression of marker genes. **(H)** Dotplot displaying sample-specific cDC2A/B gene expression in DDC D18/19 and STOP conditions. Log normalized gene expression is shown and fraction of cells expressing the genes is encoded by dot size.

This common dynamical switch towards cDC2 predominance prompted us to analyze cell state dynamics of cDC2 within our DDC atlas (cluster 5 and 15 of the main atlas, 7,563 cells), on which we performed separate clustering to obtain a more granular annotation (Fig. 3C). Clusters 3, 12, 15 were excluded from further analysis based on remnant expression of cDC1 genes. Anti-inflammatory cDC2A and pro-inflammatory cDC2B, can be distinguished based on their gene expression profiles [13]. The expression of *Clec10a* and *Mgl2*, hallmark genes for a cDC2B identity, indicated a predominant pro-inflammatory phenotype of the cDC2 subset, while maturation markers *Ccr7* (cluster 11) or *Ly6c2* (cluster 4) reflected temporal and differentiation-related changes along the x-axis in the UMAP representation (Fig. 3D). Prompted by these observations we focused on trajectory analysis within the cDC2B compartment. Disease onset was associated with the expression of “maturation-on” (MAT-ON) genes (cluster 11), that have been described in the context of DC maturation [41] and included, e.g., *Ccl17, Ccl22*, and *Arpin* (Fig. S4C). Cells derived from later timepoints (from D5 onwards) included DC precursors (preDCs) expressing *Cd7, Ly6c2, Tcf4, Runx2, Siglech*, and *Csf1r* (Fig. S4C). The majority of cDC2 derived from D9 onwards were *Mgl2*-negative, suggesting that *Mgl2* expression may be a proxy for the cDC2B maturation state.

We applied VarID2 [24] to infer transition probabilities between clusters which further supported the concept of a cDC2B trajectory linking disease-specific cell states, where *Mgl2*^+^ clusters were interconnected as part of the early DC maturation trajectory (clusters 1 – 7 – 9; Fig. S4D). To systematically investigate gene expression dynamics during maturation, we calculated *self-organizing maps* (SOMs) of pseudotemporal gene expression profiles along this trajectory (Fig. S4E) [26]. The genes in SOM modules 5, 13, and 20, which were upregulated along the trajectory (Fig. S4F) encoded soluble factors, transcription factors or proteins interacting with Nfkb1/2 (Relb, Nfkbia, Nfkbid, Nfkbiz; Fig. S3F). Since cDC2B have been linked to the induction of Il17-responses [42], we next examined the expression of genes within the gene ontology pathway “positive regulation of T helper 17 type immune responses” in the *Mgl2*^+^ clusters. Indeed, increased expression of pathway genes, such as *Jak2, Nfkbiz, Nfkbid, Nlrp3* was observed along the *Mgl2*^+^ cDC2B disease trajectory (clusters 1 – 7 – 9, Fig. S4G). Il17-response induction at the biliary niche was supported by DDC-induced *Il23a* production in BECs (Fig. 2B,C).

For the validation of increased recruitment of preDCs in prolonged DDC diet, Mgl1 and Mgl2 were co-detected using immunohistochemistry, which marks immature cDCs and macrophages [43]. Mgl1/2^+^ cells were specifically enriched in inflamed biliary foci and total numbers were increased at DDC D10 and D19 compared to controls (Fig. 3E). To assess the maturation state of hepatic cDC2, we further investigated, if the loss of *Mgl2* mRNA within cDC2 in prolonged disease can be equally observed on the protein level. Indeed, the relative proportion of Mgl2^+^ cDC2B was significantly reduced from 34.4% ± 0.2 % (D0) to 19.8% ± 4.9% at D10 (Fig. 3D, F). Finally, we tested the plasticity of Mgl2^neg^ cDC2 precursors by comparing cDC2 derived from long-term DDC diet (D18/19) to cDC2 that underwent an episode of disease recovery (DDC D18/19 + 7 days of recovery, “STOP”; Fig. 3G). Analyzing cDC2A/B gene signatures revealed induction of the restorative cDC2A gene signature during disease resolution (cluster 2 and 6), while proinflammatory cDC2B genes remained partially expressed (Fig. 3H).

In conclusion, the cDC2 scRNA-seq data reveal DC maturation and temporal exhaustion of liver-resident Mgl2^+^ cDC2B at disease progression along with recruitment of Mgl2^neg^ preDCs that may differentiate towards a proinflammatory or restorative phenotype depending on the niche signal input.

### Heterogeneous *γδ*T cells produce profibrogenic Il17a in a niche-dependent manner

To better delineate T cell subsets with key functions during the inflammatory response and tissue fibrosis, we determined *Il17a* expressing T cell populations in our scRNA-seq atlas [44]. The main *Il17a* expressing population were *γδ*T cells (cluster 15) co-expressing *Trdc, Sox13* and *Blk* [45] (Fig. S5A, B), which segregated into *Il17a*^high^ cells positive for *Scart1* (encoded by *Cd163l1*), *Trdv4* and *Trgv6*, and *Il17a*^low^ cells expressing *Scart2* (encoded by *5830411N06Rik*) (Fig. S4C). Since *γδ*T cells were sparsely represented in our atlas (282 cells) we multiplexed enriched *γδ*T cells from a hepatic control (D0), DDC D5- and D9-fed mice for scRNA-seq (Fig. 4A, B). Gene signatures defining V 6^+^ and V 4^+^ *γδ* cells indicated that Scart1^+^ cells were V 6^+^, while Scart2^+^cells were V 4^+^ [46]. Both populations expressed *Nr4a1* and *Cd5* indicating T cell receptor (TCR) activation, and *Scart1*^+^ *γδ*T17 cells (clusters 0, 3) expressed *ll17a* at higher levels (Fig. 4B, S5D). *γδ*T17 cells expressed different receptors that would enable their interaction with cDC2 and induce Il17 production, including, e.g., the Ccl2-Ccr2, Ccl20-Ccr6, Il1b-Il1r1, Il23a-Il23r, Ccl17/Ccl22-Ccr4 axes (Fig. 4C). These observations suggest a multifaceted crosstalk of *γδ*T cells with BEC and cDC2, expressing *Ccl17, Ccl22* and *Il1b* (Fig. S4C). To extend this observation, we conducted a ligand-receptor analysis of a merged liver cDC2 and *γδ*T dataset using CellChat [47] (Fig. S6A-C). This analysis further identified contact dependent and soluble interactions including Icosl-Icos, Pvr-Cd226 or Cxcl16-Cxcr6, Il18-Il18r1, Lgals9-Cd44 (Fig. S6B,C).

**Figure 4.**
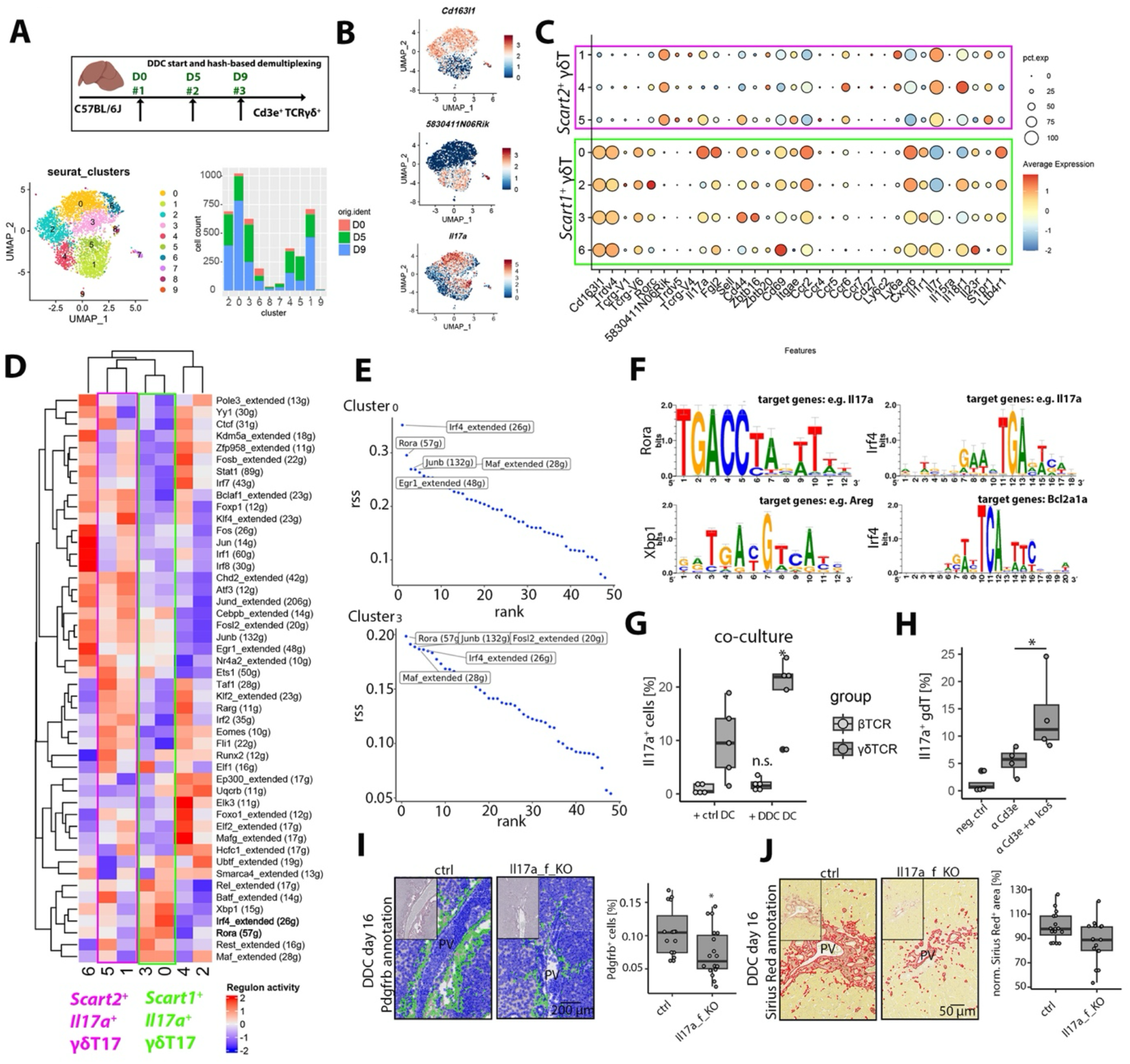
*Scart1*^*+*^ and *Scart2*^*+*^ *γδ*T cells produce profibrogenic Il17a in a niche-dependent manner. **(A)** Top: experimental design and structure of the multiplexed liver *γδ*T dataset. Bottom left: UMAP representation of the *γδ*T17 subset. Barplot displaying *γδ*T17 cluster-specific absolute cell numbers of multiplexed D0, D5, and D9. **(B)** UMAP representation of log-normalized expression of *γδ*T cell markers *Scart1* (Cd163l1), *Scart2* (5830411N06Rik) and *Il17a*. **(C)** Dotplot displaying cluster-specific gene expression. Log-normalized gene expression is color coded and fraction of cells expressing the gene is encoded by dot size. Boxes highlight *Scart1*^+^ (green) and *Scart2*^+^ (magenta) clusters. **(D)** Heatmap displaying SCENIC-inferred TF regulon activity in *Scart1*^*+*^/*Scart2*^*+*^ *γδ*T17 clusters (boxes highlight Il17a positive clusters). Predicted TF regulon activity is displayed as a color code. **(E)** Rank visualization of *Scart1*^*+*^ *γδ*T17 cluster-specific regulon specificity scores (rss). Top 5 regulons are indicated. **(F)** Predicted Rora-, Irf4- and Xbp1-specific binding motifs and target genes. **(G)** Boxplots displaying frequency of Il17a^+^ cells within Cd3^+^ *β*TCR^+^ (light grey) and Cd3^+^ *γδ*TCR^+^ (dark grey) compartments after hepatic DC coculture derived from control or DDC D5 treated mice. One-sided t-test (five independent experiments), *p ≤ 0.05; n.s., not significant. **(H)** Boxplots displaying frequency of Il17a^+^ cells in Cd3^+^ *γδ*TCR^+^ cells after anti-Cd3e or anti-Cd3e + anti-Icos stimulation. One-sided t-test (four independent experiments). *p < 0.05. **(I)** Left: Pdgfrb immunohistochemistry and positive cell detection (green overlay) of control (ctrl) and Il17a_f_KO mice after 16 days of DDC diet. Right: Boxplot displaying percentage of Pdgfrb^+^ cells. One dot represents one quantified tissue section. Two-sided t-test (n=4 mice per group). *p < 0.05. **(J)** Left: Sirius Red histochemistry and positive area detection (encircled red areas) of control (ctrl) and Il17a_f_KO mice after 16 days of DDC diet. Right: Boxplot displaying percentage of stained area. One dot represents one quantified tissue section. One-sided t-test (n=4 mice per group). *p < 0.05.

To support the hypothesis of differential signaling patterns in *Scart1*^*+*^*/Scart2*^+^ *γδ*T cells giving rise to distinct transcriptional regulation, we ran transcription factor (TF) regulon predictions using SCENIC [29]. TF regulons exhibited differential activities within and between *Scart1*^+^ and *Scart2*^+^ *γδ*T17 subsets (Fig. 4D). For example, *Scart1*^+^ *γδ*T17 cell regulons included Rora, Maf, Irf4 and Xbp1, whereas Klf2 and Klf4 were more active within *Scart2*^+^ *γδ*T17 cells, in accordance with published data [48, 49]. We then used regulon specificity scores (rss) to assess TF regulon cell type specificity. Rora, Maf, Irf4 TF regulon activity in *Scart1*^*+*^ *γδ*T17 cells (cluster 0 and 3) displayed high rss values, while Klf2 and Jund where specific for *Scart2*^*+*^ clusters 1 and 5 (Fig. 4E, S6D). Motif analysis for Rora, Irf4 and Xbp1 predicted target gene expression in *Scart1*^*+*^ *γδ*T cells including *Il17a* (Fig. 4F). Flow cytometric analysis of Il17a on *αβ* T cells (*β*TCR^+^) and *γδ*T cells confirmed increased *γδ*T17 cell numbers in cholangitis at D5/6 and D10 (Fig. S6E).

To interrogate, if *γδ*T and *αβ*T cells compete for signal cues, Tcrd-knockout (KO) mice were treated with DDC. Indeed, compensation of the *γδ*T17 function was reflected by a higher percentage of Il17a^+^ *αβ*T cells in Tcrd-KO versus control mice at DDC D5, suggesting potential competition for *γδ*T17/Th17 polarization cues in liver cholestasis (Figure S6F). Conditioned medium of DDC D5 derived DCs showed a trend towards inducing increased Il17a^+^ *γδ*T cell frequencies (p=0.06), however, compared to direct co-culture, the fraction of Il17a+ cells was low (Fig. S6G). Direct co-culture of DDC D5- or control liver-derived DCs with UTCs from control spleen led to increased Il17a production in *γδ*T cells but only weak induction in *αβ*T cells, indicating a DC-derived contribution to the induction of a *γδ*T17 cell state (Fig. 4G).

Based on this observation, we used TCR stimulation *in vitro* to further elucidate the impact of contact-based/soluble mediators in inducing a *γδ*T17 cell state. Specific stimulation with recombinant Il18, Cxcl16, Lgals9 and activating Icosl-antibodies revealed a moderate *γδ*T17 cell state induction for Icos activation, while soluble mediators did not induce significant differences (Fig. 4H, S6H). To directly test the effect of Il17a on fibrosis, we treated wildtype (wt) and Il17a/f KO mice with a long term DDC diet (16 days) and quantified the number of Pdgfrb^+^ myofibroblasts and collagen (Sirius Red) in tissue sections. The number Pdgfrb^+^ cells and collagen area was significantly reduced in Il17a/f KO versus wt mice, mirroring a profibrogenic role of Rorc-induced Il17-responses [50] (Fig. 4I,J). Similarly, tissue-derived transcript levels for *Acta2, Des, Col1a1* were reduced in DDC-treated Il17a/f KO vs controls (Fig. S6I). Localization analysis of *γδ*T cells, using a *γδ*T cell reporter mouse model (Tcrd-GDL [19]) enabled detection of *γδ*T cells in DDC-induced cholestatic livers. *γδ*T cells were detectable in lobular and portal regions, with certain *γδ*T showing a distinct peribiliary localization (Fig. S7A).

The presence of TRDC^+^, CD207^+^ cells was further verified using conventional immunohistochemistry on a human PSC collective (Fig. S7B,C). Furthermore, conducting duplex in situ hybridization on human PSC tissues, we detected niche colocalization of CLEC10A^+^/TRDC^+^ and CD207^+^/ TRDC^+^ cells (Fig. S7D).

We performed scRNA-seq analysis of two PSC patients, and conducted ligand-receptor interaction analysis for *γδ*T cells and cDC2 (Fig. S7E-G). Again, soluble and contact-based interactions were identified demonstrating a high level of cross-species conservation between cDC2 and *γδ*T cells, where 29.6% of identified interactions from mouse were found to be conserved in human (Fig. S7G, S6B,C). Using an independent PSC T cell reference [51], we tested the expression of receptors in the UTC subset of the data, to determine if these cells may sense cDC2-derived signal cues (Fig. S7H-J). Again, IL17A transcript levels could not be detected at high levels within the UTC compartment, while IL17A protein levels were detectable throughout the whole subset (Fig. S7H,I). The receptor expression overall indicated a similar molecular fingerprint within *γδ*T cells and other UTC subpopulations towards cDC2-derived cues (Fig. S7J).

Together, these results highlight Il17a-producing *γδ*T17 cells as an early-onset fibrosis-supporting communication node in murine cholangitis, while in human disease, *γδ*T cells and other UTC may have overlapping and yet undefined roles in PSC disease pathophysiology.

### Early cDC2B depletion and replacement by epigenetically restricted cDC2 results in reduced *γδ*T17 effector differentiation

To uncover the role of Mgl2^+^ cDC2B in regulating *γδ*T cell recruitment and function in the context of DDC-induced biliary injury, we depleted cDC2B using diphtheria toxin (DT) in Mgl2-DTR mice [16] (“DDC D5-DT”, Fig. 5A, S8A). The cDC2 subset of these data confirmed an almost complete abrogation of *Clec10a*^+^ *Mgl2*^+^ cDC2Bs and maturation-associated gene expression was reduced, while preDC-associated genes were up-regulated (Fig. 5B, S8B). Pathway enrichment among differentially expressed genes between cDC2 from the depleted versus control animals reflected a dysfunctional, preDC-associated cellular state according to gene signatures that affect cell migration (GPCR, chemokines), and outside-in signaling (TNFR1 signaling, Fig. S8C). Upon cDC2B depletion, BECs exhibited reduced expression of pro-inflammatory mediators *Tnf, Il23a, Ccl2* and *Ccl20* (Fig. S8D). Top differentially expressed genes within the *γδ*T cell subset included *Il17a*, which was significantly reduced in cDC2-depleted animals (Fig. 5C,D).

**Figure 5.**
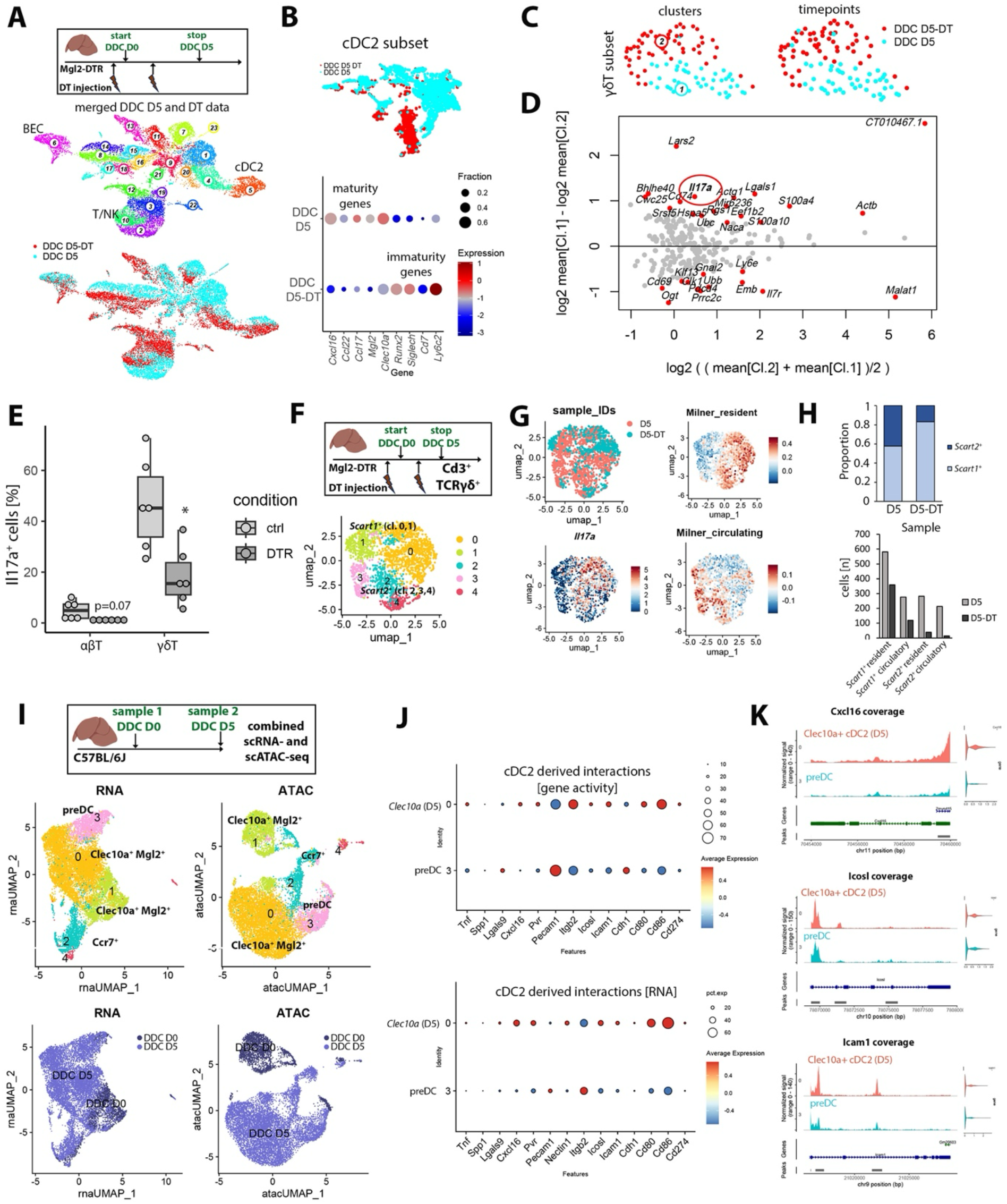
Early cDC2B depletion results in reduced *γδ*T17 effector differentiation. **(A)** Top: experimental design and structure of the merged DDC D5 and DDC D5-DT data. Bottom: UMAP representation of isolated cell types and samples. **(B)** Top: UMAP representation of cDC2 subset from data in (A) (top). Bottom: Dotplot displaying expression of immaturity and maturity markers in cDC2. Log-normalized gene expression is color coded and fraction of cells expressing the gene is encoded by dot size (bottom). **(C)** UMAP representation of *γδ*T cell subset clusters (left) and samples (right) of merged DDC D5 and DDC D5-DT data. **(D)** Differential gene expression analysis of DDC D5 and DDC D5-DT derived *γδ*T cells. Red circle highlights *Il17a*. Significantly differentially expressed genes (padj<0.05) are highlighted in red. **(E)** Boxplot displaying frequency of Il17a^+^ αβ and *γδ*T cells in DDC D5 and DDC D5-DT measured by FACS. Two-sided t-test (three independent experiments, n=6 mice per group). *p < 0.05. **(F)** Top: Experimental design of multiplexed liver *γδ*T cell experiment (DDC D5 and DDC D5-DT). Bottom: UMAP representation of *γδ*T17 clusters. **(G)** UMAP representation of multiplexed *γδ*T17 samples and log-normalized expression of *Il17a* and resident and circulatory gene signatures from [32]. **(H)** Top: stacked barplot displaying sample-specific proportions of *Scart1*^*+*^ and *Scart2*^*+*^ *γδ*T subsets in DDC D5/DDC D5-DT data. Bottom: absolute numbers of *Scart1*^*+*^ and *Scart2*^*+*^ resident and circulatory populations in DDC D5/DDC D5-DT data. **(I)** Experimental design (top). UMAP of merged control and DDC D5 derived scRNA-seq and scATAC-seq data modalities highlighting clusters derived for the data modality (middle), and sample identity (bottom). **(J)** Dotplot displaying cDC2-derived *γδ*T cell interacting gene activities (top) and RNA expression (bottom) across clusters 3 (preDC) and 0 (Clec10a^+^ cells). Log-normalized mean expression is color-coded and fraction of cells expressing the gene is encoded by dot size. **(K)** Coverage plots displaying gene-specific pseudobulk accessibility tracks in preDC (cluster 3) and Clec10a^+^ cells (cluster 0). Genomic regions of interest and coordinates are displayed in the bottom row. Violinplots on the right display RNA expression of the corresponding gene.

Flow cytometric analysis of Mgl2-depleted versus control animals revealed a significant reduction of Il17a^+^ *γδ*T cell frequencies, with a similar trend in *β*TCR^+^ *αβ*T cells (p=0.07, Fig. 5E). To capture liver *γδ*T cells at higher resolution, we enriched *γδ*T cells from Mgl2-depleted mice for scRNA-seq and interrogated the frequency distribution of *γδ*T17 cells by comparing DDC D5 and DDC D5-DT derived *γδ*T cells (Fig. 5F). Reduced total *Scart1/2*^*+*^ *γδ*T17 cell numbers were observed in the DT condition, with a more prominent reduction in the *Scart2*^+^ population (44% vs. 85% reduction, Fig. 5G,H). Further subtyping of *Scart1/2*^*+*^ *γδ*T17 cells revealed individual clusters with circulatory and resident gene signatures [31, 32], respectively (Fig. 5G, S8E,F). A reduced proportion of circulatory cells in the D5-DT versus the D5 condition indicated altered *γδ*T cell recruitment and tissue half-life in cDC2B-depleted mouse livers (Fig. 5G,H).

To address how cDC2B depletion and replacement of immature cDC2 may affect *γδ*T17 polarization across molecular layers, we performed a combined single-cell transcriptome RNA-seq and Assay for Transposase-Accessible Chromatin using sequencing (ATAC-seq) analysis (10x Multiome) on a control and a disease time-point, where all subclusters including preDCs can be observed (control (D0) and DDC D5, see methods; Fig. 5I, S9A-C). In the cDC2 subset, label transfer of RNA to ATAC clusters revealed similar manifold architecture and transitioning cell states in both data modalities proposing concordant cDC2 cell state regulation (Fig. 5I). While the Mgl2 locus did not exhibit obvious differential chromatin accessibility across cell states, the Ccr7 locus showed an increase in accessibility along the cDC2 differentiation trajectory (Fig. S9D).

This prompted us to test the genomic accessibility of identified DC-*γδ*T cell ligand-receptor pairs (Fig. S6), to screen for interactions that are epigenetically regulated and may explain reduced cDC2- *γδ*T cell interactions after mature cDC2B depletion. Multiple predicted interactions as part of the cDC2- *γδ*T cell communication node showed concordant reduced genomic accessibility and RNA expression in preDCs compared to Clec10a^+^ cDC2 (Fig 5J, S9E). Individual coverage plots of key cDC2-derived signal cues exemplified reduced genomic accessibility in preDC compared to Clec10a^+^ cDC2 (Fig. 5K, S9F).

Taken together, these data propose perturbed circulatory and residency properties of *Scart1*^+^ and *Scart2*^+^ *γδ*T17 cells and reduced *γδ*T17 effector differentiation upon cDC2B depletion, partially due to reduced genomic inaccessibility of *γδ*T cell ligands within preDCs.

### *γδ*T17 effector differentiation is induced by Mgl2^+^ cDC2B in liver draining lymph nodes

After the identification of a *γδ*T17 instructive role of cDC2B, we asked whether mature *Mgl2*^+^ *Ccr7*^+^ cDC2B induce Il17 production in liver-draining *lymph nodes* (LN), thereby promoting Th17/ *γδ*T17 responses in an organ-site independent manner [22] (Fig. 6A). To enable the identification of *Mgl2*^+^ *Ccr7*^+^ cDC2 undeterred by the maturation-associated mRNA expression loss of typical DC markers [41], we performed CITE-seq on liver-draining LN at steady state (D0), DDC D5 and D19 (19,338 cells; Fig. 6A,B). This permitted a subclassification of cDC1/2 in LN on the protein level since proteins have a much longer half-life than RNA. Indeed, the *Ccr7*^*high*^ DC cluster in the liver-draining LN dataset exhibits a clear separation into Cd11b^+^/Mgl2^+^ and Xcr1^+^ cells on the protein level, while the cDC2 subset displayed the loss of RNA marker genes (Fig. 6C, S10A). Raw numbers of CD301b^+^ cDC2Bs in the process of DDC-associated inflammation continuously increased in the LN scRNA-seq data mirroring liver egress (Fig. 6B-D). In liver draining LN, Mgl2^+^ cDC2 were in physical contact with *γδ*T cells, as seen in Tcrd-GDL mice (Fig. 6E). Contact and ligand-receptor-mediated interaction analysis confirmed the previously observed DC- *γδ*T cell signaling module (Fig. 6F,G). Again, multiple liver interactions including Icosl-Icos, Cxcl16-Cxcr6, Icam1-Itgal/Spn were also predicted in LN promoting the idea of site-independent cDC2- *γδ*T cell interactions in cholestatic liver disease. Next, we analysed a scRNA-seq dataset containing *γδ*T17 cells derived from liver draining

**Figure 6.**
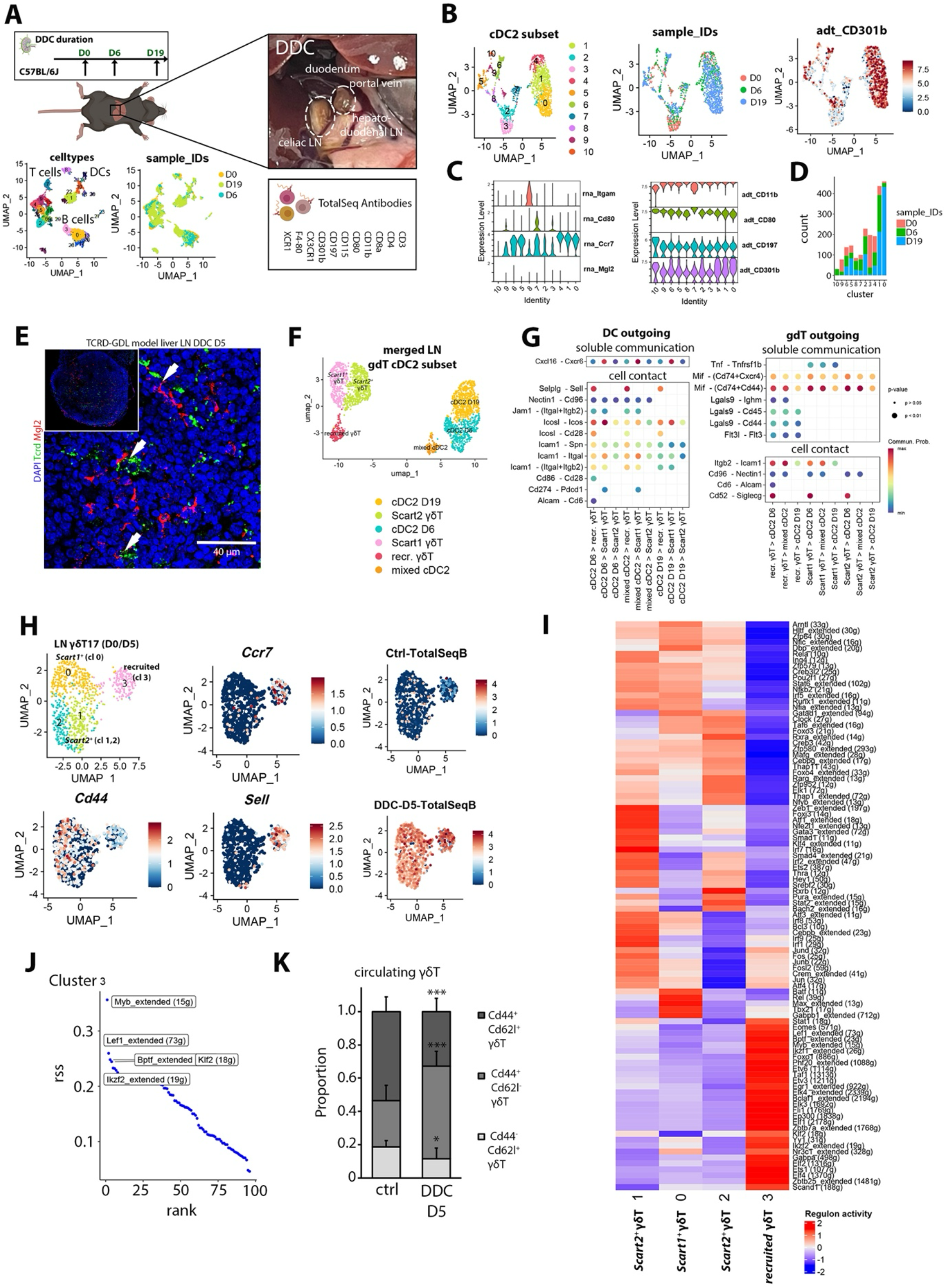
*γδ*T17 polarization is induced by Mgl2^+^ cDC2B in liver draining lymph nodes. **(A)** Experimental design of the merged liver draining LN control, DDC D6 and DDC D19 CITE-seq dataset. Macroscopic view of the hepatoduodenal and celiac trunk lymph nodes in situ [22]. **(B)** UMAP representation of the LN-derived cDC2 clusters (left), timepoints (right) and centered log-ratio transformed CD301b (encoded by *Mgl2*) protein expression. **(C)** Violinplots displaying log-normalized RNA expression for *Itgam, Cd80, Ccr7* and *Mgl2* (left) and centered log-ratio transformed protein expression for the corresponding proteins (right). **(D)** Barplot displaying timepoint-specific absolute cell numbers of cDC2 in LN. **(E)** IF displaying DAPI, Mgl2 co-stained liver-draining LN of DDC D5-treated Tcrd-GDL mice (n=3). Scale bar, 40 µm. **(F)** UMAP representation of merged LN *γδ*T cDC2 subset displaying clusters used for cell communication inference. **(G)** Dotplot displaying cDC2 and *γδ*T cell outgoing molecular contact-based and soluble communication pairs. Communication probability encoded as color code and p-values as dot size. **(H)** UMAP representations displaying clusters of LN *γδ*T17 cells derived from DDC D0/D5 timepoints, log-normalized *Ccr7, Sell, Cd44* expression and centered log-ratio transformed data of sample hashtags. **(I)** Heatmap displaying SCENIC-inferred TF regulon activity in *Scart1*^*+*^ (cluster 0), *Scart2*^*+*^ (clusters 1,2) and recruited (cluster 3) *γδ*T17 clusters shown in (H). Predicted TF regulon activity is displayed as a color code. **(J)** Rank visualization of recruited *γδ*T17 cluster-specific regulon specificity scores (rss). Top 5 regulons are indicated. **(K)** Stacked barplot displaying relative proportions of Cd44^+^ Cd62l^+^ (central memory-like), Cd44^+^ Cd62l^−^ (effector) and Cd44^−^ Cd62l^+^ (naïve) circulating *γδ*T cell populations. Two-sided t-test (3 independent experiments, n=7-8 mice/group). *p < 0.05, ***p < 0.001.

LN of D0 and D5 and detected *Ccr7*^*+*^ *Sell*^*+*^ *Lck*^+^ (cluster 3), *Scart1*^+^ (cluster 0) and *Scart2*^+^ (clusters 1,2) *γδ*T17-polarized cells. We performed SCENIC analysis to identify TF networks uniquely associated with these LN *γδ*T17 cells (Fig. 6H-J). A unique cassette of TF-regulons with predicted activity restricted to *Ccr7*^*+*^ *Sell*^*+*^ *γδ*T17 cells (“recruited”, cluster 3) was identified, with the highest rss for the TFs Myb, Lef1, Bptf and Ikzf2, in accordance with the RNA expression levels (Fig. 6H, S10B). Altered *γδ*T17 frequencies in LN prompted us to analyze *γδ*T in the circulation. Indeed, FACS analysis of circulating *γδ*T cells from ctrl (D0) and DDC D5 timepoints revealed altered *γδ*T dynamics, with a decrease of naïve Cd62l^+^ Cd44^−^ and Cd62l^+^ Cd44^+^ circulatory *γδ*T cells, while Cd44^+^ Cd62l^−^ effector *γδ*T cells increased (Fig. 6K). Last, we compared *γδ*T17 cells derived from draining LN at DDC D5 and upon loss of *Mgl2*^+^ cDC2B at DDC D5-DT. *γδ*T17 populations derived from the D5-DT were almost completely abrogated in our scRNA-seq data, further indicating a connection of LN *γδ*T17 effector differentiation and cDC2 homing (Fig. S11A-C).

Altogether, we here characterized a communication network between cDC2 and *γδ*T cells, where altered cDC2B dynamics impact on *γδ*T17 effector differentiation in liver and draining LN, exerting a profibrogenic function in liver cholangitis.

## Discussion

While a number of immunological susceptibility genes have been identified in PSC [52], the disease-aggravating and self-perpetuating biliary niche-specific immunological circuits that maintain and drive disease pathology remain incompletely understood.

Il17 has been identified as a key driver of liver disease and fibrosis [53], yet *γδ*T cell-derived Il17a as an important cellular source has been widely neglected. While our data hints towards a profibrogenic role of *γδ*T cell-derived Il17a, *γδ*T cell-specific knockout models will be required to fully elucidate the role of these cells in liver scarring.

To understand the heterotypic interplay of the biliary niche including cDC2 interacting with *γδ*T cells, we here generated a DDC liver and LN-specific single-cell reference. Local and distant *γδ*T17 responses were mediated by *Scart1*^*+*^ and *Scart2*^*+*^ *γδ*T cell subpopulations, which have been described to exhibit both resident and circulating properties. Although antigen processing and presentation by MHCs are thought to be dispensable for *γδ*T cell activation [54, 55], our results show that antigen-presenting cDC2B within the immunobiliary niche may contribute to a *γδ*T17 response, e.g. through contact-dependent signals such as Icosl-Icos interaction. While these results need to be confirmed with independent and possibly more cell-specific depletion models, our data imply that Icosl and additional ligands derived from cDC2 may promote *γδ*T17 cell states, while expression of these ligands in cDC2 was partially regulated by genomic accessibility.

The here inferred TF regulon activity extends known concepts of divergent *γδ*T17 populations and their transcriptional regulation in disease [56]. While *γδ*T cell landscapes in multi-organ settings have revealed site-dependent states [33], our study highlights functional *γδ*T responses in primary organ and the tissue-draining LN. For example, the chemokine receptor Ccr6 known to be essential for *αβ*T and *γδ*T positioning in liver fibrosis [57, 58] was expressed in *Scart2*^+^ but not *Scart1*^+^ *γδ*T cells. *Scart2*^+^ cells also express *S1pr1* and *S1pr2* for which opposing functions for regulating tissue residency and egression have been described in skin *γδ*T cells [59, 60]. Our data imply a role for *Scart2*^*+*^ *γδ*T cell recruitment in cholangitis, while resident *Scart1*^+^ *γδ*T cells, were the main Il17a producers. Despite these phenotypic differences, *γδ*T17 cells showed a dependence on cDC2B across subtypes.

The presence of naïve *Sell*^*+*^ *Ccr7*^*+*^ *γδ*T cells in LN at steady state and early disease (D5) may indicate, that *γδ*T cell niche replenishment from the circulation depends on restorative signal cues, as these cells contribute to tissue regeneration and scarring [61].

Context-dependent signal input from neighboring cells as part of cellular crosstalk may affect cDC2B tissue retention and *γδ*T17 differentiation in liver and LN. The biliary tree contains a unique immunological niche [62], and our DDC and PSC data corroborate such multi-dimensional molecular crosstalk through spatial and *in silico* validation of immunobiliary myeloid (e.g. via Csf1, Ccl2, Mif, Lgals9) and *γδ*T cell (e.g. via Il18, Cxcl16, Icosl) interactions. However, UTC composition and heterogeneity may not be fully conserved across species [63]. Nonetheless, our scRNA-seq analysis of published data [51] indicates a comparable expression signature of cDC2-interacting genes across UTC subtypes, thereby supporting the functional unit-concept of UTC and partial transferability of the cDC2- *γδ*T cell interactions from mouse to human [64].

Moreover, *γδ*T cell accumulation was detectable in different mouse models of cholestatic liver injury (DDC, BDL), indicating a more general disease response. More profound analysis of all *γδ*T cell states, spatial and multimodal analyses are required to get a more holistic understanding about potential disease-specific mechanisms. In that regard, our DDC atlas also captures naïve *γδ*T, *γδ*T1 cells, and cellular dynamics of other disease promoting myeloid cells, which should be further explored [10, 65, 66].

Altogether, our site- and time-resolved cholangitis atlas provides novel insights into heterocellular interactions of underexplored immune cells at unprecedented resolution and captures the dynamics of hepatic cDC2 and *γδ*T cells within the immunobiliary niche.

## Abbreviations

BEC, biliary epithelial cell; cDC, conventional dendritic cell; CEC, continuous endothelial cell; DDC, 3,5-Diethoxycarbonyl-1,4-Dihydrocollidine Diet; DR, ductular reaction; EC, endothelial cell; *γδ*T, gamma delta T cell; GSEA, gene set enrichment analysis; HE, hematoxylin eosin; KC, Kupffer cell; LSEC – liver sinusoidal endothelial cell; LN, lymph node; PSC – primary sclerosing cholangitis; SD, standard deviation; TMA, tissue microarray

## Author Contributions

D.G. and S.T. conceived and coordinated the project. S.T. designed and performed experiments and computational analysis. H.H. and S.B. supported FACS and co-culture analysis. H.H. performed D.C. in vitro experiments. A.A. supported bioinformatic analysis. J.S. provided support with mouse strains. S. provided support for the design and analysis of T cell experiments. F.I. provided support with sequencing. Ta.P. contributed immunohistochemistry experiments. M.T., A.R. provided PSC histological assessment. K.B.H., C.E.Z. supported human liver tissue analysis. To.P., J.K. provided processed human PSC scRNA-seq data. N.R. supervised K.B.H.. D.G. supervised the project. S.T. and D.G. wrote the manuscript. All authors read and edited the manuscript.

## Acknowledgements

The authors thank Konrad Schuldes, Sebastian Hobitz, Nadine Vornberger, Rémi Doucet Ladevèze, Wiebke Brethauer, Tim Rau, Carolin Kerber for excellent technical assistance. We thank Akiko Iwasaki and Kristin Hogquist for providing Mgl2-DTR mice, and Wolfgang Kastenmüller and Martin Väth for providing Il17a/f-KO, Tcrdtm_KO and Tcrd-GDL mice. We would like to thank the Core Unit for FACS of the IZKF Würzburg and Christian Linden for supporting this study. We would like to thank the Center of Model System and Comparative Pathology (CMCP) and Christine Schmitt, Heike Conrad, Diana Lutz, Karin Rebholz for their support. We would like to thank Sabine Roth from the Institute of Pathology in Würzburg.

We thank Haristi Gaitantzi, Vanessa Hartwig (Department of Surgery, Medical Faculty Mannheim), Emrullah Birgin (Department of Surgery, University Hospital Ulm) and Chang-Feng Chu (Leibniz Institute for Natural Product Research and Infection Biology, Jena) for their support with fresh human liver tissue. We would like to thank the Single-Cell Center Würzburg, and the Core Unit Systems Medicine and Panagiota Arampatzi, Mugdha Srivastava, Pierre Khoueiry and Tom Graefenhahn. We would like to thank the Deep Sequencing Facility at the Max Planck Institute for Immunobiology and Epigenetics (Chiara Bella and Ulrike Böhnisch) and Helmholtz Centre for Infection Research Braunschweig (Michael Jarek, Maren Scharfe and Doris Järke). We would like to thank Heike Wagner and Sabine Kranz, Stefan Brandt, Gallina Miller, Kim Patricia Rohr and Denice Schmitt (Animal Facility of the Würzburg Institute of Systems Immunology). We want to thank Martin Väth, Christin Friedrich and Milas Ugur for helpful discussions. Schemes within manuscript figures were generated with BioRender.

