## Supplementary Figures for "An immunobiliary single-cell atlas resolves crosstalk of type 2 cDCs and γδT cells in cholangitis"

[1]Würzburg Institute of Systems Immunology, Julius-Maximilians-Universität Würzburg, Würzburg, Germany
[2]University Hospital Freiburg, Department of Gastroenterology, Hepatology, Endocrinology and Infectious Diseases, Freiburg, Germany
[3]Helmholtz Institute for RNA-based Infection Research (HIRI), Helmholtz-Center for Infection Research (HZI), Würzburg, Germany
[4]Center for Model System and Comparative Pathology, Institute of Pathology, University Hospital Heidelberg, Heidelberg, Germany
[5]University Hospital Heidelberg, Institute of Pathology, Heidelberg, Germany
[6]Leibniz Institute for Natural Product Research and Infection Biology, Jena, Germany & Friedrich Schiller University Jena, Germany
[7]I. Department of Medicine, University Medical Center Hamburg-Eppendorf, Hamburg, Germany
[8]current address: Cancer Immunology Program, Peter MacCallum Cancer Centre, Melbourne, VIC, Australia; Sir Peter MacCallum Department of Oncology, The University of Melbourne, Melbourne, VIC, Australia
[9]Center for Immunology, University of Minnesota, Minneapolis, Minnesota, USA
[10]Institute of Pathology, University of Würzburg, Würzburg, Germany
[11]Department of Surgery, Medical Faculty Mannheim, Heidelberg University, Mannheim, Germany
[12] Department of Surgery, University Hospital Ulm, Ulm, Germany
[13] CAIDAS - Center for Artificial Intelligence and Data Science, Würzburg, Germany

### Figure S1

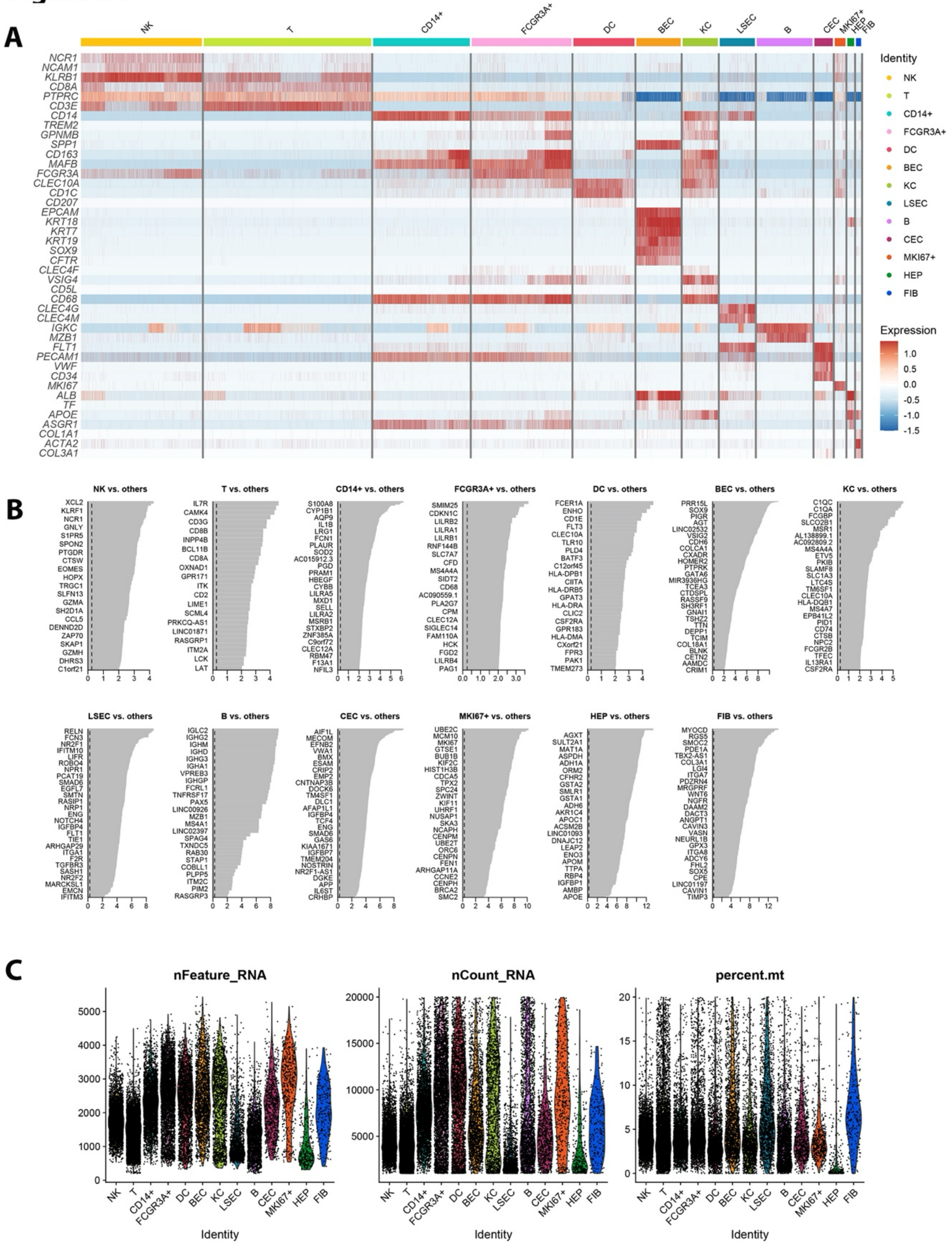


**Figure S1. Characterization of human scRNA-seq data**

**(A)** Heatmap displaying log-normalized marker gene expression for all cell types of the human single cell data reference (n=6 patients). **(B)** Barplots displaying 25 upregulated cluster-specific genes (cluster of interest vs. all other clusters). **(C)** Violinplots displaying cluster-specific QC-metrics of the human scRNA-seq dataset.

### Figure S2

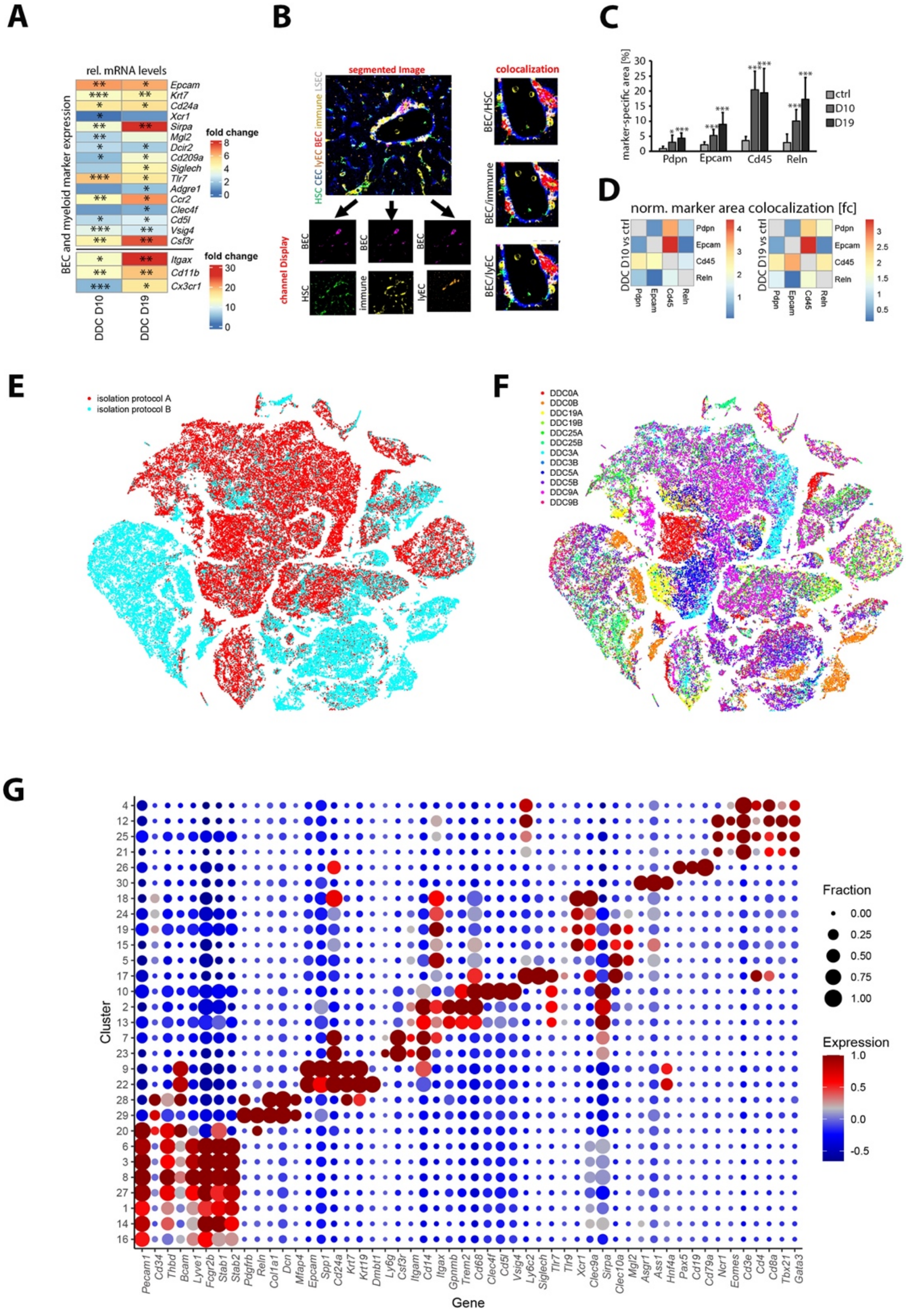


**Figure S2. Immunobiliary niche in DDC-treated mice and DDC scRNA-seq data**

**(A)** Channel-specific display of IF of the immunobiliary niche on steady state mouse liver tissue (compare main Fig. 1E). **(B)** Heatmap displaying relative mRNA levels (qPCR) of cell type marker genes in DDC D10/D19 liver tissues normalized to liver control. Two sided t-test (n=4 mice/group), *p < 0.05, **p < 0.01, ***p < 0.001. **(C)** Graph displaying IF-inferred channel-specific co-localization from intensity-based segmentation in the portal niche. Region of interest (right) highlights colocalized marker areas in red. **(D)** Marker-specific area quantification of segmented IF images derived from control (ctrl), DDC D10 and D19 liver tissues (Fig. 1E,H). Two-sided t-test (n=3-4 mice/group; 3 images per mouse), *p ≤ 0.05, ***p ≤ 0.001. **(E)** Heatmap displaying average colocalization foldchange (fc) within different color channels in DDC D10, D19 liver tissues normalized to control. **(F)** tSNE representation of the cell composition of the DDC atlas data based on isolation protocol A/B (Methods). **(G)** tSNE representation of the independent samples of the DDC atlas data. A/B indicates the isolation strategy of the respective timepoint (Methods). **(H)** Dotplot displaying candidate genes for the DDC single cell atlas reference for clusters with at least 200 cells. Color code encodes for Z score of the mean expression across clusters and dot size represents fractions of cells being positive.

#### Figure S3

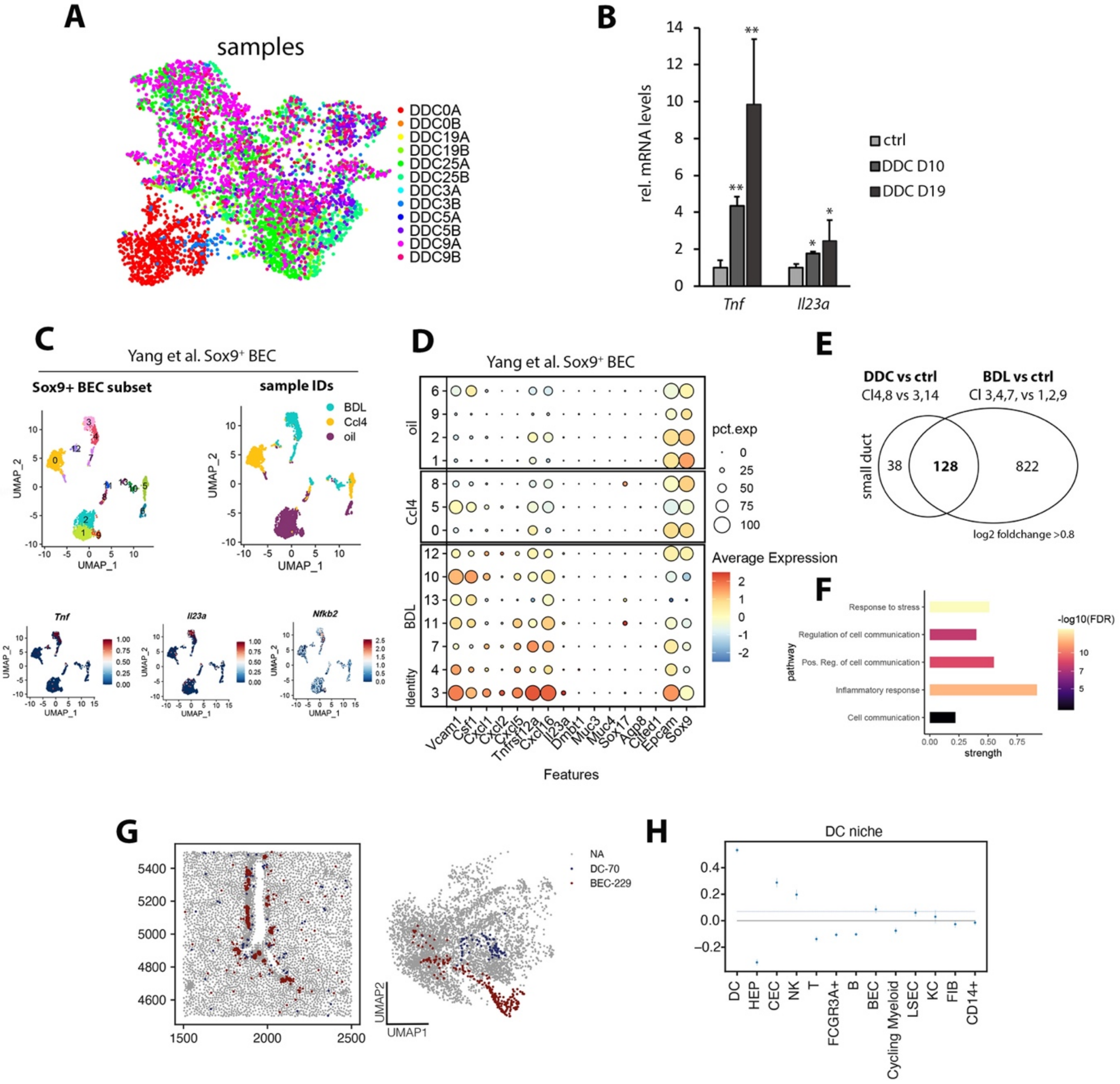


**Figure S3. DDC induces ductular reaction and small duct inflammatory disease**

**(A)** UMAP representation of sample-specific labeling of the BEC subset. **(B)** Barplot displaying relative *Tnf* and *Il23a* mRNA levels (qPCR) in DDC D10/D19 liver tissues normalized to liver control. Asterisks indicate level of significance. Two-sided t-test (n=4 mice/group), $^*p < 0.05$, $^{**}p < 0.01$. **(C)** UMAP of Sox9⁺ BEC clusters (left) and treatment conditions (right) of BDL and CCl4 treated mice. “Oil” represents respective control. UMAP of log-normalized expression of *Tnf*, *Il23a*, *Nfkb2*. Data derived from (Yang et al., 2021). **(D)** Dotplot displaying BEC cluster-specific gene expression of the BDL Sox9⁺ data subset derived from (Yang et al., 2021). Log-normalized average expression and fraction of cells expressing gene of interest shown. BDL derived cluster 3 was identified as Sox9+ BEC cluster with pro-inflammatory changes. **(E)** Venn diagram displaying commonly upregulated genes (log2-foldchange > 0.8) in Sox9⁺ BDL and DDC data. **(F)** Pathway enrichment analysis of the common BDL/DDC-upregulated genes. **(G)** Region of interest (same as in Figure 1A) displaying colocalized DC and BEC niche. **(H)** Logistic regression coefficients of the DC niche. Regression coeffcients (y-axis) were ordered by magnitude. Grey dotted line indicates threshold (c=0.07) for plotting the cell type niche interaction map in Figure 2F. Error bars indicate SD from five cross fold trainings of logistic regression.

### Figure S4

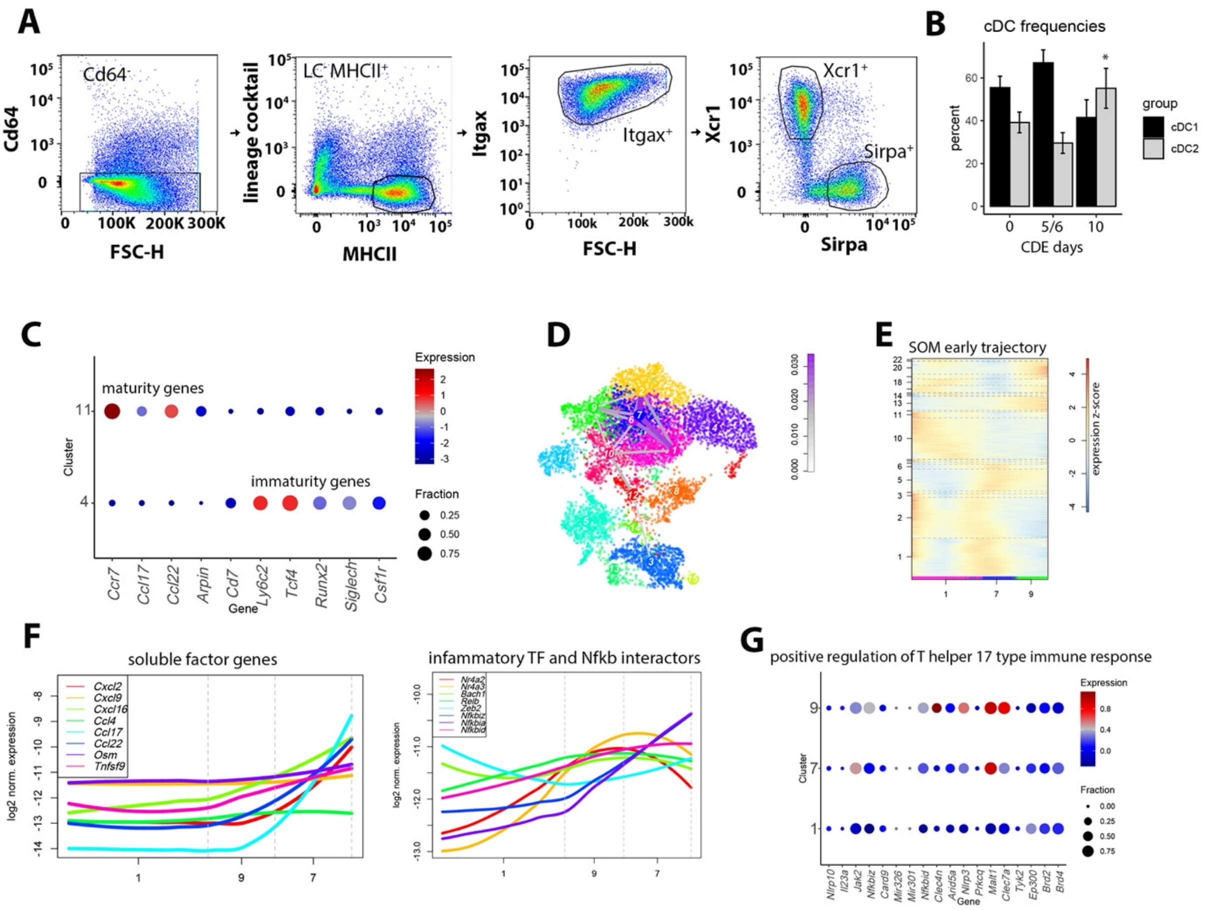


**Figure S4. Disease stage-specific cell state transition of cDC2 reflects immature niche restoration**

(**A**) FACS gating strategy used to quantify cDC1 and cDC2 derived from [1]. **(B)** Barplot displaying cDC1/cDC2 FACS-based quantification at D0, CDE D5/6 and D10. One-sided Wilcoxon test (n=3-4 mice per group, two independent experiments) *$p < 0.05$. Control datapoints (“0”) derive from Fig. 3B. **(C)** Dotplot displaying cDC2-specific gene expression of maturation associated genes. Log-transformed gene expression is color coded and fraction of cells expressing the gene is encoded by dot size. **(D)** UMAP of cDC2 subset overlaid with VarID2-derived transition probabilities. **(E)** Pseudotime-ordered self-organizing map (SOM) of the inferred early disease trajectory (cluster 1 – 7 – 9). **(F)** Pseudotemporal gene expression profiles of representative soluble factor (left) or proinflammatory and Nfkb interacting genes (right). The sum of gene expression across all cells was normalized to one. **(G)** Dotplot displaying cDC2 cluster-specific gene expression with the annotation “positive regulation of T helper 17 type immune response”. Z score of the mean expression is color coded and fraction of cells expressing the gene is encoded by dot size.

### Figure S5

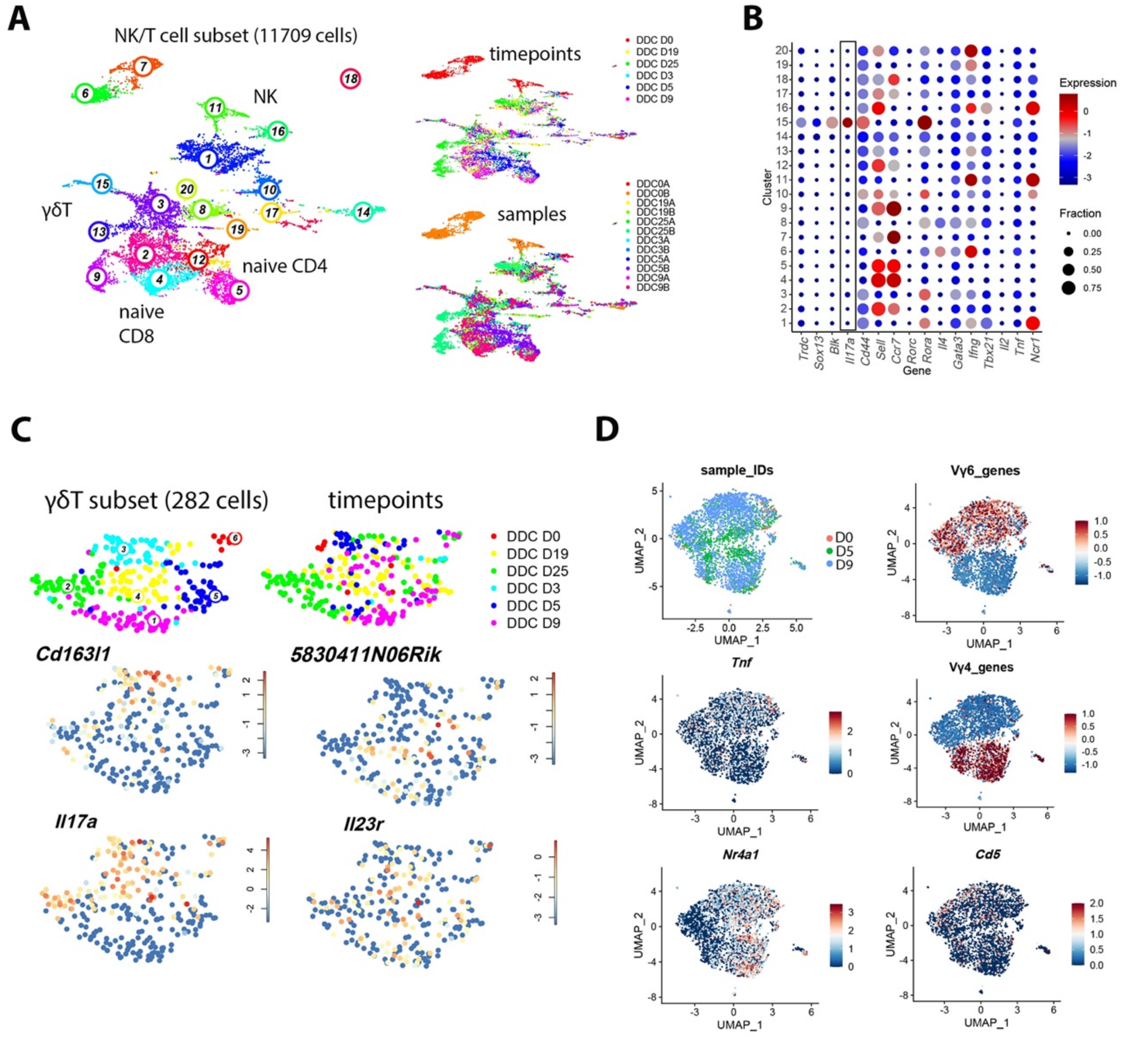


**Figure S5. *Scart1⁺* and *Scart2⁺* γδT cells produce profibrogenic Il17a in a niche-dependent manner**

**(A)** UMAP of DDC T/NK cell subset (11,709 cells) with cluster, timepoint and sample annotation. **(B)** Dotplot displaying NK/T cell cluster-specific gene expression. Log-normalized gene expression is color-coded and fraction of cells expressing the gene is encoded by dot size. Box highlights *Il17a*, which is exclusively expressed in cluster 15. **(C)** UMAP representation of γδT cell subset (282 cells) with cluster annotation and individual timepoints (top). Log-normalized expression of *Scart1* (Cd163l1), *Scart2* (5830411N06Rik), *Il17a* and *Il23r*. **(D)** UMAP representation displaying timepoint-specific data distribution in the γδT17 subset (top left). Log-normalized expression of Vγ6/Vγ4 genes and *Tnf, Nr4a1, Cd5* in γδT17 cells.

#### Figure S6

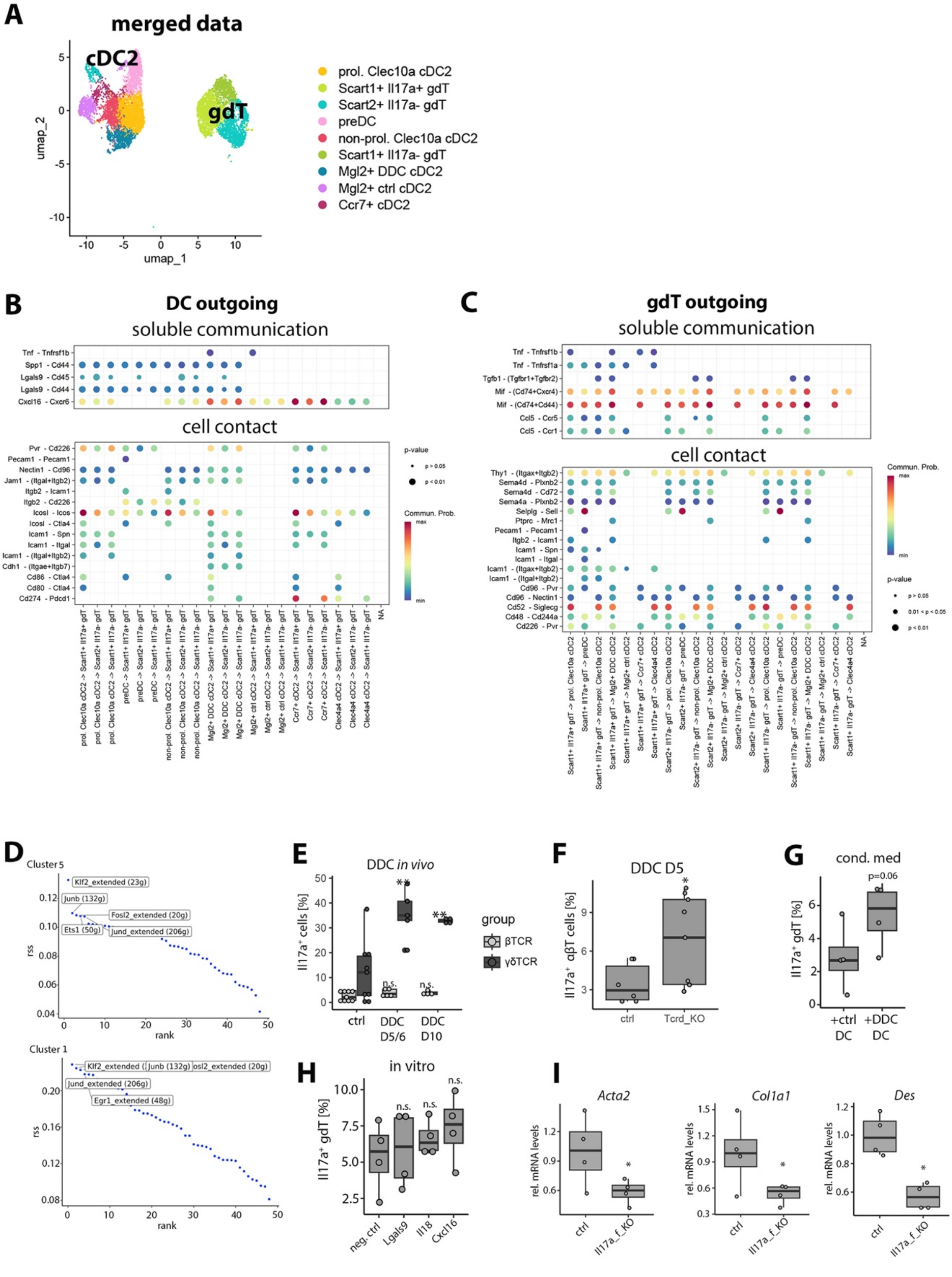
A
merged data
cDC2
gdT
umap_1
umap_2
prol. Clec10a cDC2
Scart1+ Il17a+ gdT
Scart2+ Il17a- gdT
preDC
non-prol. Clec10a cDC2
Scart1+ Il17a- gdT
Mgl2+ DDC cDC2
Mgl2+ ctrl cDC2
Ccr7+ cDC2
B
DC outgoing
soluble communication
cell contact
C
gdT outgoing
soluble communication
cell contact
D
Cluster 5
Cluster 1
rank
rss
E
DDC in vivo
Il17a⁺ cells [%]
ctrl
DDC D5/6
DDC D10
group
βTCR
γδTCR
F
DDC D5
Il17a⁺ αβT cells [%]
ctrl
Tcrd_KO
G
cond. med
p=0.06
Il17a⁺ gdT [%]
+ctrl DC
+DDC DC
H
in vitro
Il17a⁺ gdT [%]
neg. ctrl
Lgals9
Il18
Cxcl16
I
Acta2
Col1a1
Des
rel. mRNA levels
ctrl
Il17a_f_KO

**Figure S6. γδT cell communication and profibrogenic, niche-dependent Il17a production**

**(A)** UMAP representation of merged liver γδT and cDC2 cell subsets displaying clusters used for cell communication inference. **(B)** Dotplot displaying cDC2 outgoing molecular cell contact and soluble communication pairs. Communication probability encoded as color code and p-values as dot size. **(C)** Dotplot displaying gdT outgoing molecular cell contact and soluble communication pairs. Communication probability encoded as color code and p-values as dot size. **(D)** Rank visualization of *Scart2*$^{+}$ γδT17 cluster-specific regulon specificity scores (rss). Top 5 regulons are indicated. **(E)** Boxplots displaying frequency of Il17a$^{+}$ cells within Cd3$^{+}$ βTCR$^{+}$ (light grey) and Cd3$^{+}$ γδTCR$^{+}$ (dark grey) compartments after cytokine re-stimulation. Two-sided t-test (4 independent experiments, n=5-9 mice/group). *$p < 0.05$, **$p < 0.01$; n.s., not significant. **(F)** Boxplots displaying proportion of Il17a$^{+}$ cells within Cd3e$^{+}$ βTCR$^{+}$ cells after cytokine re-stimulation in control (ctrl) and Tcrd_KO mice. Two-sided Wilcoxon test (three independent experiments, n=6-9 mice/group) *$p < 0.05$. **(G)** Boxplots displaying frequency of Il17a$^{+}$ cells within Cd3$^{+}$ γδTCR$^{+}$ cells after in vitro administration of ctrl and DDC D5 DC-derived conditioned medium. One-sided t-test (4 independent experiments). **(H)** Boxplots displaying frequency of Il17a$^{+}$ cells within Cd3$^{+}$ γδTCR$^{+}$ cells after in vitro administration of Lgals9, Il18, Cxcl16. One-sided t-test (4 independent experiments). n.s., not significant. **(I)** Boxplots displaying relative *Acta2, Col1a1* and *Des* mRNA levels (qPCR) in DDC D16 treated ctrl and Il17a_f_KO liver tissues normalized to liver control. Asterisks indicate level of significance. One-sided t-test (n=4 mice/group), *$p < 0.05$.

### Figure S7

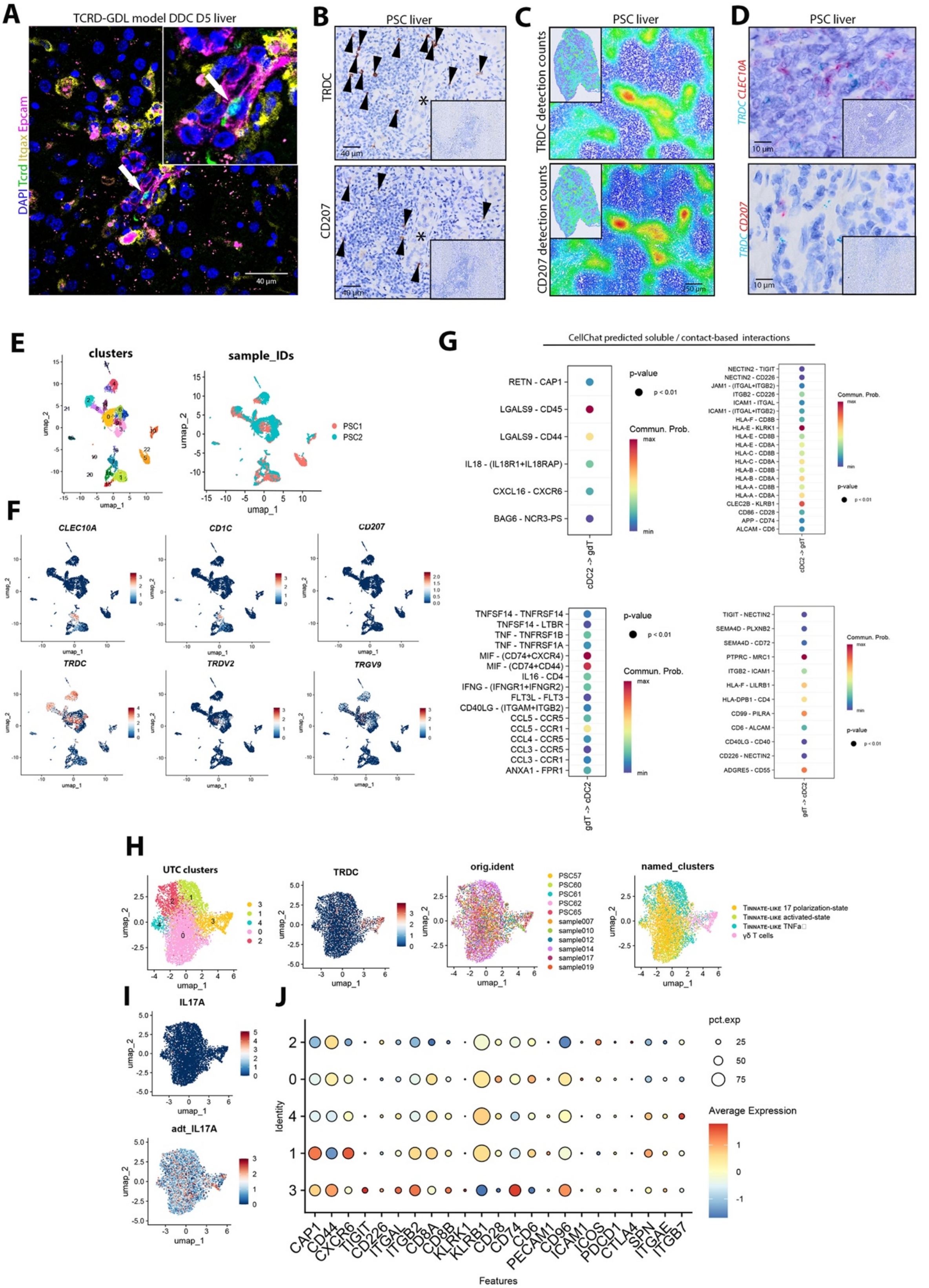

A
TCRD-GDL model DDC D5 liver
DAPI Tcrd Itgax Epcam
40 µm
B
PSC liver
TRDC
CD207
40 µm
C
PSC liver
TRDC detection counts
CD207 detection counts
D
PSC liver
TRDC CLEC10A
TRDC CD207
10 µm
E
clusters
sample_IDs
PSC1
PSC2
umap_1
umap_2
F
CLEC10A
CD1C
CD207
TRDC
TRDV2
TRGV9
G
CellChat predicted soluble / contact-based interactions
RETN - CAP1
LGALS9 - CD45
LGALS9 - CD44
IL18 - (IL18R1+IL18RAP)
CXCL16 - CXCR6
BAG6 - NCR3-PS
cDC2 -> gdT
p-value
p < 0.01
Commun. Prob.
max
min
TNFSF14 - TNFRSF14
TNFSF14 - LTBR
TNF - TNFRSF1B
TNF - TNFRSF1A
MIF - (CD74+CXCR4)
MIF - (CD74+CD44)
IL16 - CD4
IFNG - (IFNGR1+IFNGR2)
FLT3L - FLT3
CD40LG - (ITGAM+ITGB2)
CCL5 - CCR5
CCL5 - CCR1
CCL4 - CCR5
CCL3 - CCR5
CCL3 - CCR1
ANXA1 - FPR1
gdT -> cDC2
H
UTC clusters
TRDC
orig.ident
named_clusters
PSC57
PSC60
PSC61
PSC62
PSC65
sample007
sample010
sample012
sample014
sample017
sample019
I
IL17A
adt_IL17A
J
pct.exp
25
50
75
Average Expression
Identity
CAP1
CD44
CXCR6
TIGIT
CD226
ITGAL
ITGB2
CD8A
CD8B
KLRK1
KLRB1
CD28
CD74
CD6
PECAM1
CD96
ICAM1
ICOS
PDCD1
CTLA4
SPN
ITGAE
ITGB7
Features

**Figure S7. cDC2-$\gamma\delta$T cell communication in human PSC**

**(A)** IF displaying DAPI, Mgl2 co-stained DDC D5 liver derived from TCRD-GDL mice (n=3). Arrow indicates peribiliary localized $GFP^+$ cell. Scale bar, 40 µm. **(B)** TRDC and CD207 immunohistochemical staining on human PSC liver tissue (n=18). Peribiliary immune infiltrates are enriched for $TRDC^+$ and $CD207^+$ cells. Asterisk indicates bile duct. Scale bar, 40 µm. **(C)** High- and low-resolution visualization of color-coded $TRDC^+$ and $CD207^+$ detection counts in human PSC tissue. Analysis indicates varying cell concentration within different tissue compartments. Scale bar, 250 µm. **(D)** Duplex TRDC/CLEC10A and TRDC/CD207 in-situ hybridization of human PSC liver tissue (n=5). Cellular colocalization and/or close proximity relationships between cells can be detected. **(E)** UMAP representations of clusters and human PSC samples. **(F)** UMAP representations of log-normalized expression of cDC2 and $\gamma\delta$T cell marker genes. **(G)** Dotplot displaying cDC2 and $\gamma\delta$T cell outgoing molecular cell contact and soluble communication pairs. Communication probability encoded as color code and p-values as dot size. **(H)** UMAP representation of UTC subset derived from Poch et al. [2]. Original cell annotation, patient identity and TRDC expression are shown. **(I)** UMAP representation of log-normalized expression of IL17A and centered log ratio transformed IL17A protein expression. **(J)** Dotplot displaying UTC cluster-specific gene expression of cDC2-sensing receptors and interacting genes. Log-normalized average expression and fraction of cells expressing gene of interest shown.

#### Figure S8

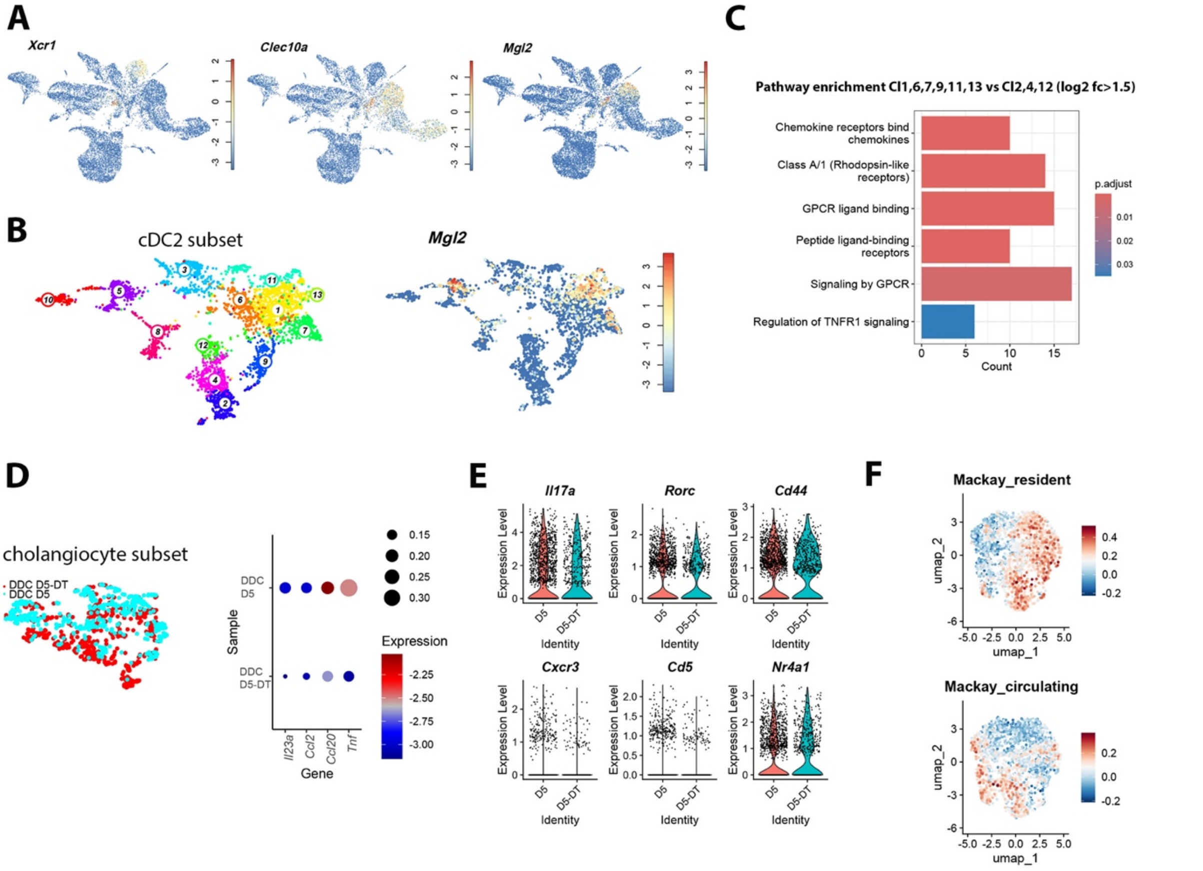


**Figure S8. Early cDC2B depletion results in reduced γδT17 effector differentiation**

**(A)** UMAP representation of the combined DDC D5/DDC D5-DT dataset displaying log-normalized gene expression of *Xcr1, Clec10a, Mgl2*. **(B)** UMAP representation displaying clusters of the cDC2 subset of merged DDC D5 and DDC D5-DT data (left). UMAP representation of cDC2 subset displaying log-normalized *Mgl2* expression (right). **(C)** Barplot displaying pathway enrichment of differentially upregulated genes in cDC2 in DDC D5 (clusters 1,3,6,7,9,11,13) vs. DDC D5-DT (clusters 2,4,12). Enriched pathways in the non-depleted condition show an association with cDC2 function. **(D)** UMAP representation of BEC subset of merged DDC D5 and DDC D5-DT data (left). Dotplot displaying expression of cholestasis associated inflammatory mediators. Log-normalized expression is color coded and fraction of cells expressing the gene is encoded by dot size (right). **(E)** Violinplots displaying log-normalized expression if Il17a, Rorc, Cd44, Cxcr3, Cd5, Nr4a1 within gdT17 subset across samples (compare Fig. 5F). **(F)** UMAP representation displaying log-normalized gene expression of Mackay resident and circulatory gene signatures in γδT17 subset derived from experiment of Fig. 5F.

### Figure S9

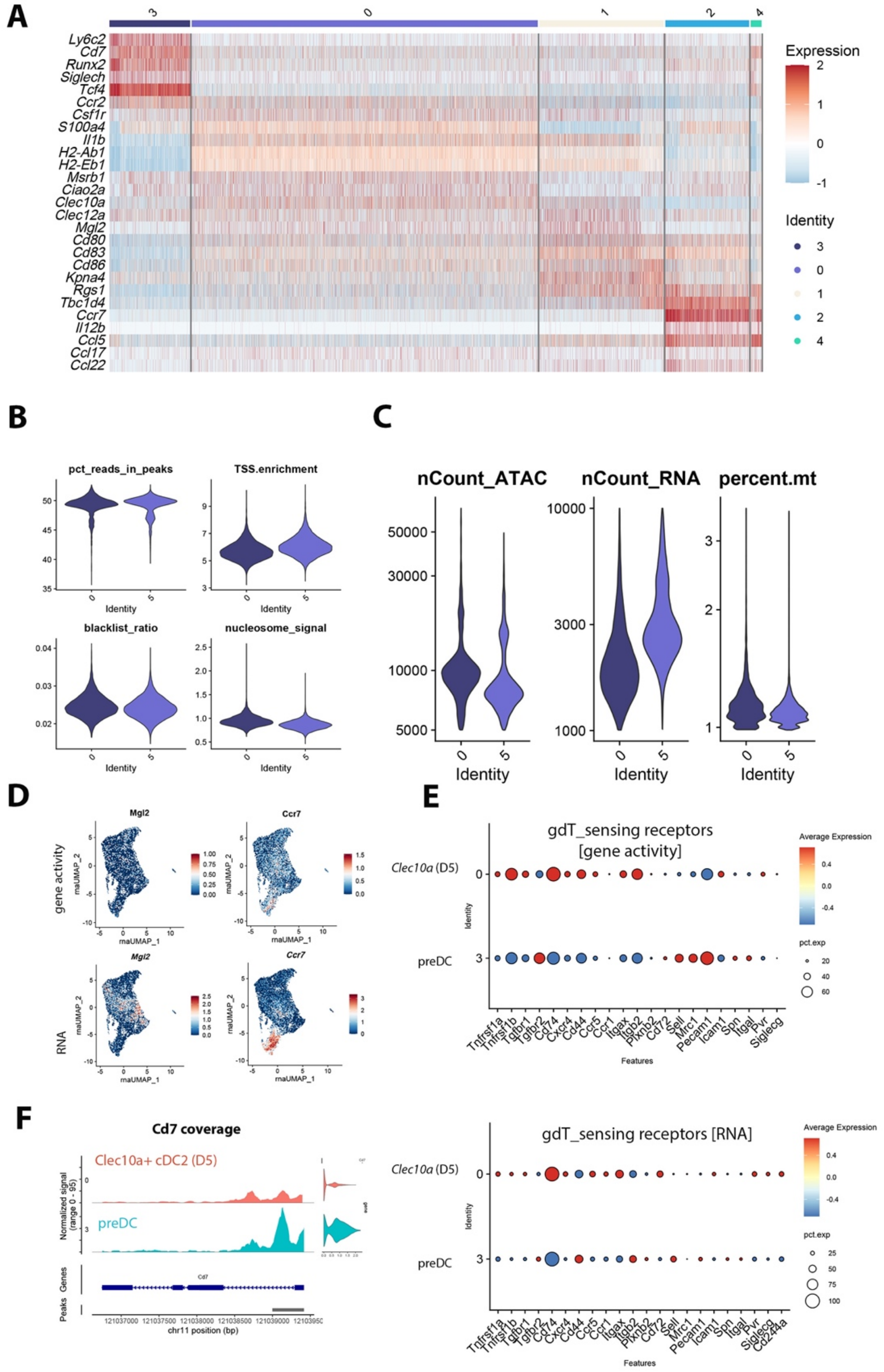


**Figure S9. Quality control metrics of Multiome data**

**(A)** Heatmap displaying log-normalized marker gene expression of the cDC2 subset data (compare with Figure 5I). **(B)** Violin plots displaying QC metrics of the ATAC-seq data derived from the DDC D0/D5 Multiome dataset. **(C)** Violin plots displaying captured ATAC/RNA counts and fraction of reads mapped to the mitochondrial genome. **(D)** scRNA-derived UMAP (rnaUMAP) highlighting gene activity levels inferred from scATAC data (top) and *Mgl2* and *Ccr7* RNA expression (bottom). **(E)** Dotplot displaying $\gamma\delta$T cell sensing receptor gene activities in cDC2 across clusters 3 (preDC) and 0 (Clec10a⁺ cells). Log-normalized mean expression is color coded and fraction of cells expressing the gene is encoded by dot size. **(F)** Cd7 coverage plot displaying gene-specific pseudobulk accessibility tracks in preDC (cluster 3) and Clec10a⁺ cells (cluster 0). Genomic region of interest and coordinates displayed in the bottom row. Violinplot on the right displays *Cd7* RNA expression.

### Figure S10

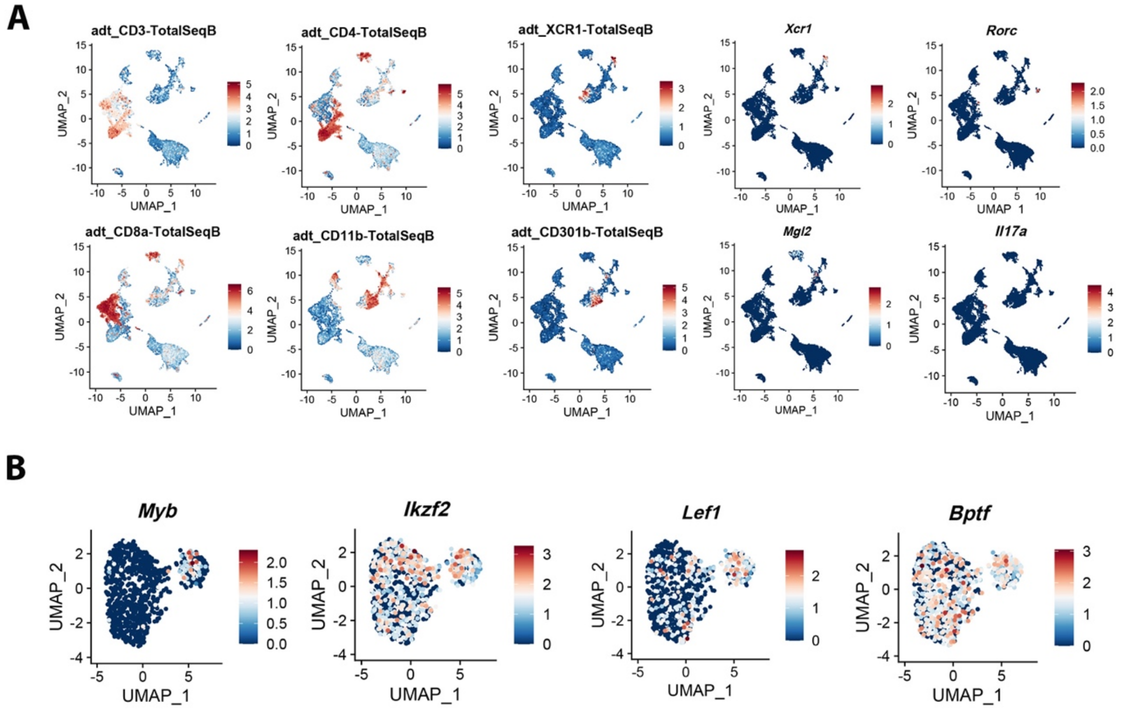


**Figure S10. $\gamma\delta$T17 polarization is induced by CD301b$^+$ cDC2B in liver draining lymph nodes**

**(A)** UMAP of the complete LN dataset displaying centered log-ratio normalized protein data (CD3, CD4, CD8, CD11b, XCR1, CD301b) and log-normalized RNA data (*Xcr1*, *Mgl2*, *Rorc*, *Il17a*). **(B)** UMAP representation displaying log-normalized *Myb, Lef1, Ikzf2, Bptf* expression.

### Figure S11

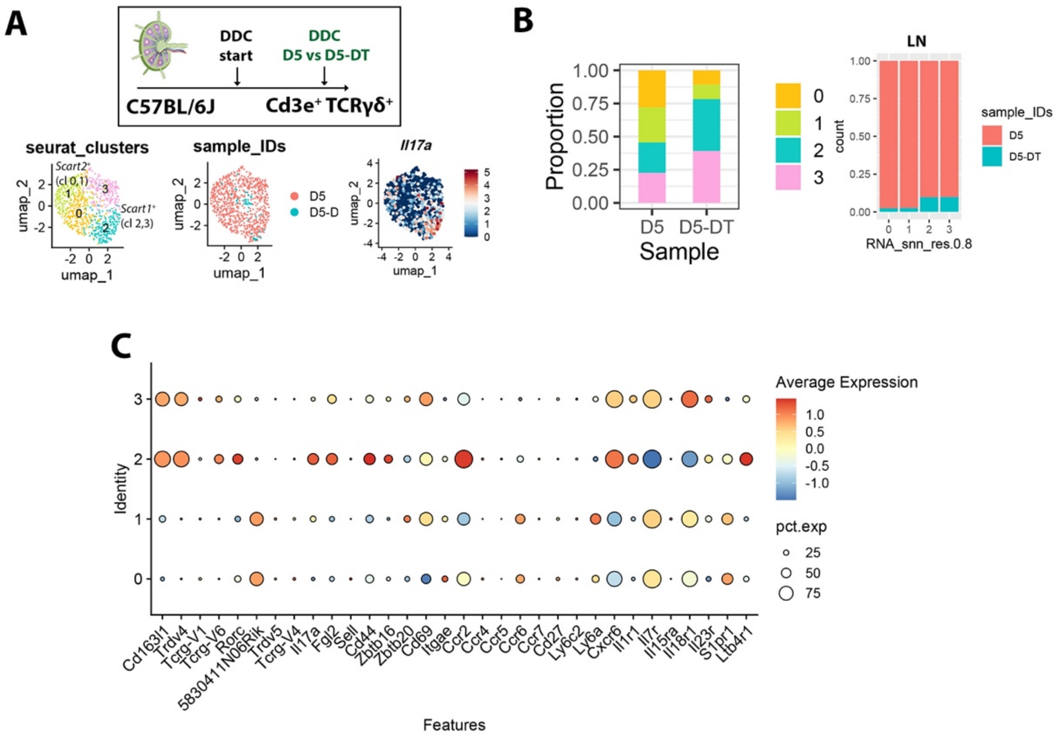


**Figure S11. CD301b⁺ cDC2B cell depletion abrogates γδT17 cell states in liver draining LN**

**(A)** Experimental design and structure of the merged liver draining LN DDC D5 and LN DDC D5-DT dataset (top). UMAP representations of clusters, samples and log-normalized Il17a expression (bottom). **(B)** Stacked barplot displaying relative proportions of γδT17 per cluster (left) and condition (right). **(C)** Dotplot displaying cluster-specific gene expression in γδT cells. Log-normalized average expression and fraction of cells expressing gene of interest shown.

#### Supplemental Information

#### Materials and Methods

**Contact**

**Data and Code Availability Statement**
scRNA-seq data will be deposited at GEO and will be publicly available as of the date of publication. Code to reproduce major results will be deposited at GitHub and will be publicly available as of the date of publication. Any additional information required to reanalyze the data reported in this paper is available from the lead contact upon reasonable request. Due to patient data protection and institutional guidelines, human tissue microarray data cannot be made publicly available.

**Cell enrichment strategies for individual scRNA-seq datasets:**

- **Liver steady state and DDC-dataset**: in two independent rounds of sequencing the following cell types were enriched at equal ratios at day 0 (ctrl), D3, D5, D9, D18 or D19, D25. For STOP and D5-DT data, only protocol B was used. D18 and D19 timepoints were summarised as D19 to facilitate timepoint-specific interpretation within the atlas. The following cell populations were sorted: 1) alive singlet, $Cd45^+$, $Lyve\text{-}1^+$, $Pdpn^+$, $Epcam^+$, $Cd31^+$ $Lyve\text{-}1^-$, retinoid-autofluorescent cells (405 nm excitation, 450/40 emission) [3]. 2) alive cells from HSC prep, $Cd45^+$ $Itgax^+$ $Cd64^-$, $Epcam^+$, $Cd3e^+$ $Lyve\text{-}1^-$.
- **Mouse CITE-seq LN dataset (ctrl and DDC):** the following cell types were enriched from pooled hepatoduodenal and celiac LNs at D0, D6, and D19 and mixed at equal ratios: alive singlet, $Cd45^+$ $Itgax^-$, $Cd45^+$ $Itgax^+$, $Pdpn^+$, $Cd45^-$ $Pdpn^-$.
- **γδT cell datasets**: cells from liver and LN were labeled using hashtag antibodies and alive $Cd45^+$ $Cd3e^+$ $TCR\beta^-$ $TCR\gamma\delta^+$ cells were sorted at indicated disease timepoints. The relative yield of $\gamma\delta$T cells in every timepoint represents the fraction of sorted cell numbers per mouse and timepoint.
- **Human liver dataset:** Pseudonormal and far distant human liver tissue was used for cell isolation from 6 patients (4 female, 2 male, age range 45-76 yrs, receiving a lobectomy or hemihepatectomy for oncologic resection) and the following cell populations were FACS enriched at equal distribution: 1) alive, 2) $CD45^+$ $CD3E^-$, 3) $CD45^+$ $CD3E^+$ 4)$CD45^+$ $CD11C^+$ 5) $CLEC4G^+$ 6) $CLEC4G^-$ $CD31^+$, 7) $EPCAM^+$. This gating strategy was applied to enrich for immune cells of the immunobiliary niche, while the specific localization of the respective cells was not further assessed. Cell types of the human atlas were clustered at low resolution, so that LAM did not cluster separately, but were part within the $FCGR3^+$ cluster. Lymphatic endothelial cells and portal fibroblasts as important cell types of the portal/immunobiliary niche were not actively enriched for in our sorting strategy and are thus not part of the dataset.
- **Human liver PSC dataset:** Liver explant tissues from two patients with end-stage PSC were used for cell isolations (patient 1, 53 yrs, female: Ustekinumab amongst others; patient 2, 21 yrs, male: Mesalazine and Vedolizumab amongst others). The following cell populations were FACS enriched at equal distribution: 1) alive, 2) $CD45^+$, 3) $CD3E^+$ $TCRgd^+$ 4) $CD3E^+$ $TCRgd^-$ 5) $CD45^-$ 6) $CD11C^+$ $CD64^+$ 7) $CD11C^+$ $CD64^-$.

**Histochemical and immunohistochemical analysis of human and mouse liver tissues**

3 µm thick sections of a previously characterized human *tissue microarray* (TMA) with a core size of 1.5 mm that contained 78 pseudonormal liver tissues [4] was immunohistochemically stained for CK7, CD34, PDPN, CD3, TRDC and CD207. Antigen retrieval was performed using Ventana CC1 solution and primary antibodies were incubated 24 min – 40 min. DAB solution was added after linker- and multi-incubation and sections counterstained with haematoxylin. All used antibodies can be found in the CTAT Methods Table. TMA slides were scanned using a Pannoramic Scan II scanning device (3DHISTECH, Budapest, Hungary) with a 40x magnification. Positive cell detection of liver tissues was performed within the in-built TMA dearrayer of QuPath after setting a single intensity threshold [5]. Human PSC liver tissues (n=18) were stained against TRDC and CD207. RNAscope (Bio-Techne, Minneapolis, USA) duplex in situ hybridization of human PSC tissues was performed using commercially available probes against TRDC, CLEC10A and CD207 according to the manufacturer's recommendations. Tissues were provided in accordance with the regulations of the Tissue Bank of the National Center for Tumor Diseases (NCT) Heidelberg and the ethics committee of Heidelberg University (S-230/20, S-206, 207/05). The use of the fresh human resection material was approved under the ethics vote 2012-293N-MA and 2021-2320-1. For

immunohistochemical detection of mouse Mgl1/2, liver tissue antigen retrieval was performed at pH6 in a steamer for 8 min, and Mgl1/2 antibody (#AF4297, R&D systems, Minneapolis, USA) incubated for 1 h at room temperature (1:75 dilution). Horse anti-goat IgG coupled alkaline phosphatase-based detection was performed for 7 min. For the detection of Pdgfrb on mouse liver tissues, sections were incubated one hour with the primary antibody after heat-induced antigen retrieval at pH9. An anti-rabbit secondary antibody conjugated to AP was applied (Polyview Plus AP reagent, ENZO Life Sciences GmbH, Lörrach, Germany) and the signal was visualized using Permanent AP Red (Zytovision GmbH, Bremerhaven, Germany).

**Dendritic and T cell isolation, FACS, cultivation and semi-quantitative PCR**

Cells were isolated from 6 – 12-week-old C57Bl/6J mice. After tissue digestion and gradient centrifugation, cells were enriched using antibody-coated microbeads and LS columns (MiltenyiBiotec, Bergisch Gladbach, Germany). A published DC gating strategy was used to detect and quantify cDC1 and cDC2 [1]. For *in vitro* cytokine re-stimulation, T cells were stimulated for 3 hrs with 50 ng/ml PMA (Merck, Darmstadt, Germany), 1 µg/ml ionomycin (Merck) and 1 µg/ml Brefeldin A (Merck). After re-stimulation, cells were surface stained, fixed and permeabilized using FOXP3/Transcription Factor Staining Buffer Set (eBioscience, San Diego, USA) and stained against Il17a and data acquired using a FACSCelesta (BD, New Jersey, USA). For liver DC – splenic T cell coculture, hepatic DCs were isolated and magnetically enriched using Cd11c microbeads and LS columns according to the manufacturer's recommendations. Simultaneous to hepatic DC isolation, splenic T cells were isolated in a two-step isolation procedure to enrich the fraction of γδT cells. First, a negative selection against $CD4^+$ and $CD8^+$ conventional T cells was performed using CD4/CD8 (TIL) microbeads and LD columns (MiltenyiBiotec). The negative fraction was then enriched using CD3e microbeads and LS columns. Liver DCs and splenic T cells were seeded in a 1:1 ratio in RPMI160 (Gibco) supplemented with 10% FCS (Corning, New York, USA) and penicillin/ streptomycin (Thermo Scientific, Waltham, USA) in addition with mouse recombinant Il23 (10 ng/µl, Thermo Scientific). For TCR stimulation, 24 wells were coated over night with anti-Cd3e antibody (5 µg/ml, Biolegend), washed 3 times with PBS and enriched splenic T cells cultured for 20 hrs with Lgals9, Il18, Il23a, Cxcl16, anti-mouse Icos-activating antibody or DC-conditioned medium derived from ctrl or DDC D5 derived liver DCs (mixed with RPMI containing 10% FCS and Pen/Strep in a 1:1 ratio).

Cells were harvested for FACS analysis 20 h after culture. For the analysis of circulating γδT cells, mouse blood was withdrawn by intracardiac puncture and erythrocytes were lysed using ACK lysis buffer (Thermo Scientific) prior to FACS staining and data acquired using a Cytek Aurora, (Cytek Biosciences, Fremont, USA).

For bulk RNA isolation from cells and tissues, the Arcturus PicoPure RNA isolation kit was used (Thermo Scientific, Waltham, USA). After cDNA synthesis using RevertAid First Strand cDNA Synthesis kit (Thermo Scientific), semiquantitative PCR was set up using SYBR Green (Bio-Rad, Hercules, USA) using 40 amplification cycles as previously described [6]. Melting curves were generated to test for product amplification specificity and samples were measured in technical triplicates. Gene expression was normalized using the normaqPCR package and the measurement of three independent housekeepers (Actin, Gapdh, Hprt) [7]. Used primers are listed in the CTAT Methods Table.

**Sample preparation and fluorescence microscopy and quantification**

Snap-frozen liver tissues were cut into 5 µm thick tissue sections using a Leica CM 3050 S cryostat and fixed in ice-cold methanol. To block unspecific binding, tissue sections were incubated with goat-serum prior to incubation with primary and secondary antibodies. Sections were washed and mounted into Fluoromount G (Southern Biotech, Birmingham, USA) with or without DAPI. Six color immunofluorescence microscopy images were obtained using a Zeiss LSM780 laser scanning confocal microscope. Detectors were adjusted to detect 6 wavelength windows using 4 excitation laser wavelengths (405, 488, 561, 639 nm). The far-red laser was used for simultaneous excitation of AF639 and AF700 fluorescent dyes, which could be detected using wavelength-specific detection windows. The 405 nm laser was used for a combined excitation of BV421 and BV605. Image tiles were obtained using an oil immersion 40x objective and stitched based on a default overlap region of 10%. Four color IF images were obtained using a Olympus IX83 microscope containing a Yokogawa spinning disc unit. To optimize image display, raw images were background subtracted (rolling ball, default settings) and an adjustment of the display window performed using FIJI [8]. All images within one experiment were adjusted identically. Colocalization analysis of image channels and interaction areas were processed using the "math" function in FIJI. Average cell colocalization areas per group were calculated and displayed as a heatmap.

#### Quantification and statistical analysis

For quantification and statistical analysis, R studio was used, and data is presented as mean ± SD. Graphs were plotted using R studio or Excel and panels organized using Adobe Photoshop and Adobe Illustrator. For information regarding independent repeats of experiments, please refer to the respective figure legends. Statistical tests are specified in the figure legends. Shapiro-Wilk test was used to test for normality. Wilcoxon rank-sum test was used for non-normal distributed data. T-test was used for normally distributed data. The number of animals per group in each experiment are indicated in the figure legends. P values are indicated as following: *p≤0.05, **p≤0.01, ***p≤0.001, n.s. not significant.

#### Data and Code availability

Single-cell RNA-seq data have been deposited at GEO. Accession numbers will be publicly available as of the date of publication. For information regarding reagents, antibodies, mouse strains please refer to the separate Supplementary Methods Table.
